# Exploiting WEE1 kinase activity as FUS::DDIT3-dependent therapeutic vulnerability in myxoid liposarcoma

**DOI:** 10.1101/2024.03.13.584771

**Authors:** Lorena Heinst, Kwang Seok Lee, Ruth Berthold, Ilka Isfort, Svenja Wosnig, Anna Kuntze, Susanne Hafner, Bianca Altvater, Claudia Rössig, Pierre Åman, Eva Wardelmann, Claudia Scholl, Wolfgang Hartmann, Stefan Fröhling, Marcel Trautmann

## Abstract

The pathognomonic FUS::DDIT3 fusion protein drives myxoid liposarcoma (MLS) tumorigenesis via aberrant transcriptional activation of oncogenic signaling. Since FUS::DDIT3 has so far not been pharmacologically tractable to selectively target MLS cells, this study investigated the functional role of the cell cycle regulator WEE1 as novel FUS::DDIT3-dependent therapeutic vulnerability in MLS. Here we demonstrate that enhanced WEE1 pathway activity represents a hallmark of FUS::DDIT3-expressing cell lines as well as MLS tissue specimens and that WEE1 is required for MLS cellular survival *in vitro* and *in vivo*. Pharmacologic inhibition of WEE1 activity results in DNA damage accumulation and cell cycle progression forcing cells to undergo apoptotic cell death. In addition, our results uncover FUS::DDIT3-dependent WEE1 expression as an oncogenic survival mechanism to tolerate high proliferation and resulting replication stress in MLS. Fusion protein-driven G1/S cell cycle checkpoint deregulation via overactive Cyclin E/CDK2 complexes thereby contributes to enhanced WEE1 inhibitor sensitivity in MLS. These findings identify WEE1-mediated replication stress tolerance as molecular vulnerability in FUS::DDIT3-driven MLS tumorigenesis that could represent a novel target for therapeutic intervention.

## Introduction

Myxoid liposarcomas (MLS) are malignant soft tissue tumors of adipocytic origin accounting for 20-30% of all liposarcomas (1). They frequently arise in younger adults and represent the most common liposarcoma subtype in patients below the age of 20 years, with primary lesions predominantly occurring in the proximal extremities (2, 3). High rates of local recurrence and metastatic spread to distant sites in approximately 40% of patients contribute to MLS pathology (4–6). Morphologically, MLS encompass a spectrum ranging from paucicellular myxoid tumors to hypercellular round cell high-grade sarcomas associated with a more aggressive clinical course (7). Current treatment options comprise surgical excision and (neo-)adjuvant radiation and/or conventional chemotherapy to potentiate long-term survival in MLS patients (8). Although MLS demonstrates enhanced chemosensitivity, advanced or metastatic disease often translates into poor patient prognosis and therapy is administered with palliative intent (9). This illustrates the unmet clinical need for novel biology-guided therapeutic strategies in the treatment of MLS.

The identification of novel targets represents a challenge in MLS, since MLS, similar to other translocation-associated sarcomas, harbor few genetic alterations beyond their characteristic chromosomal t(12;16)(q13;p11) translocation juxtaposing fractions of the *FUS* gene to the entire coding sequence of *DDIT3*. The resulting FUS::DDIT3 fusion protein drives MLS tumorigenesis via sustained transcriptional activation of oncogenic signaling (10–13). Despite recent advances, the particular mode of action of FUS::DDIT3 still remains incompletely understood. Since molecular targeting of FUS::DDIT3 is notoriously challenging, the identification of novel FUS::DDIT3-essential effectors and signaling pathways represents the most promising strategy to interfere with fusion protein-driven oncogenesis and selectively target MLS cells. In this study, we therefore investigated the functional requirement for the cell cycle regulator WEE1 in MLS, which was identified as FUS::DDIT3-essential effector in a functional genomic screen in an established MLS model system.

WEE1 is a key modulator of the highly conserved replication stress response pathway. Upon activation, WEE1 mediates cell cycle arrest on intra-S and G2/M cell cycle checkpoints via inhibitory phosphorylation on tyrosine 15 on CDK1 and CDK2, complexed with cyclin A, B or E (14–16). WEE1 thereby enables ordered DNA replication in S phase and G2 cell cycle transition while attenuating replication stress and paving the way for DNA damage repair prior to mitotic entry (17–19). Tumor cells particularly depend on this function, as replication stress represents a hallmark feature of cancer (20, 21). Mechanistic contributions to replication stress are diverse, however, defective G1/S cell cycle checkpoint signaling, observed in the majority of tumor cells, is often involved (22). As compensation, affected cells rely on replication stress pathways and a functional G2/M cell cycle checkpoint to maintain genomic integrity and cellular survival (23, 24), simultaneously offering a targetable vulnerability (25–28). Consistently, recent pharmacologic approaches targeting WEE1 kinase, among them the potent inhibitor MK-1775, have shown promising efficacy (29, 30). WEE1 kinase inhibition causes cell cycle checkpoint abrogation, leads to S phase defects and forces cells with accumulated DNA damage to undergo unscheduled mitosis, which ultimately results in cell death – an unfavorable state termed mitotic catastrophe (31, 32).

Here, we evaluated the functional relevance of WEE1 in FUS::DDIT3-expressing cells by investigating the antitumor effects of pharmacologic WEE1 inhibition and the underlying mechanism for enhanced requirement for WEE1 pathway activity.

## Results

### Enhanced WEE1 expression and pathway activity in FUS::DDIT3-expressing cells

To identify novel pharmacologically addressable targets for therapeutic intervention in the biological context of the MLS-specific FUS::DDIT3 fusion protein, we first integrated the data obtained in a loss-of-function shRNA screen, which consisted of approximately 27,500 shRNAs targeting over 5,000 human genes, in immortalized mesenchymal stem cells either stably expressing the pathognomonic *FUS::DDIT3* fusion gene or an empty control vector (Figure 1A) as previously published (33). With respect to FUS::DDIT3-selective essentiality, we identified WEE1, a central G2/M cell cycle checkpoint regulator, among the top kinases with high essentiality score in FUS::DDIT3-expressing SCP-1 cells (Figure 1B). Subsequent GSEA pathway analysis demonstrated a high frequency of genes involved in the cell cycle machinery, particularly mitosis and DNA replication, among the ten most essential pathways in FUS::DDIT3-expressing SCP-1 cells (Figure 1C).

**Figure 1.**
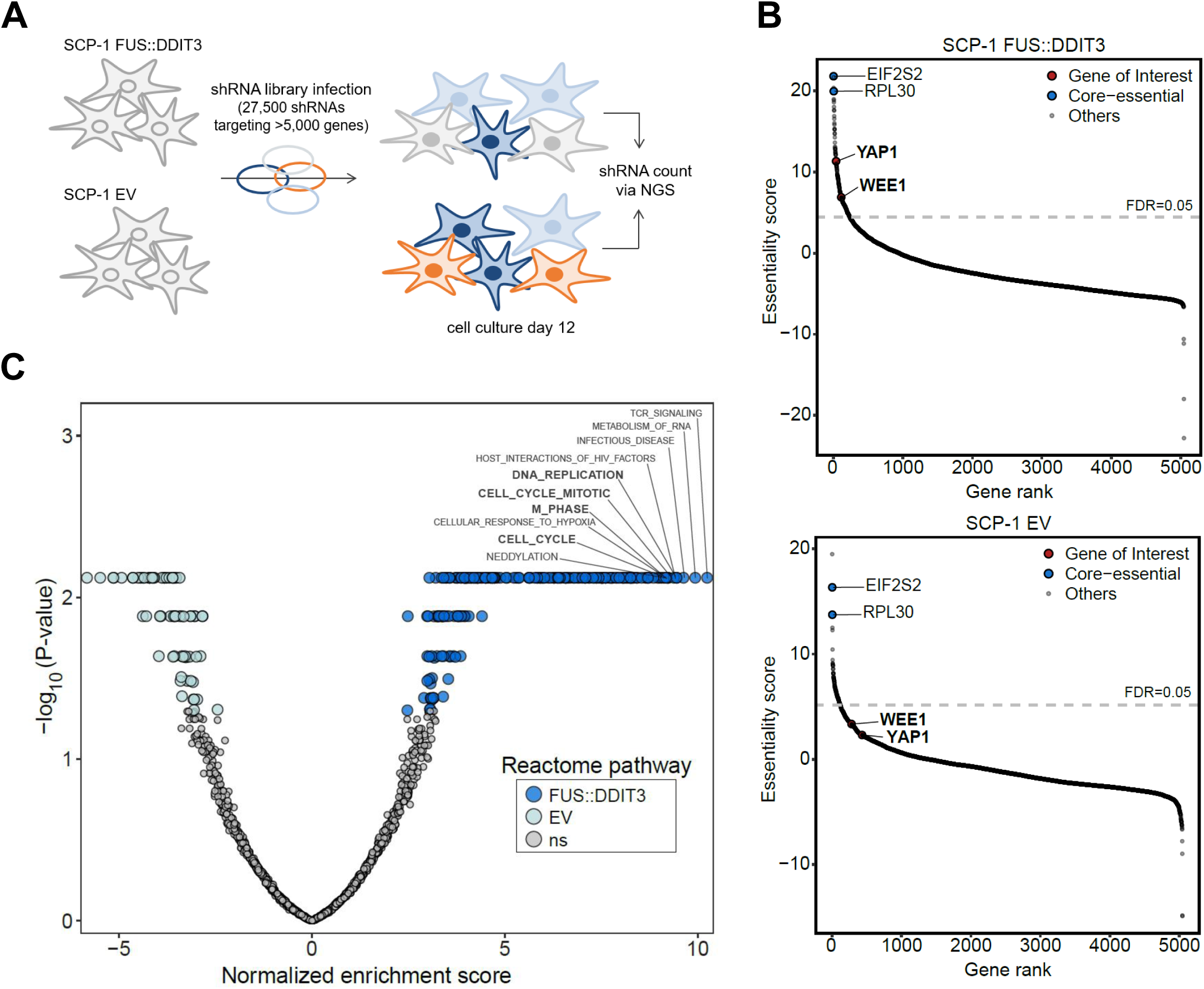
Identification of WEE1 kinase as requirement in FUS::DDIT3-expressing mesenchymal stem cells. **A** Schematic illustration of RNAi-based loss-of-function screen conducted in mesenchymal stem cells (SCP-1) expressing the chimeric *FUS::DDIT3* fusion transcript or an empty control vector (EV) (33). **B** Genes were ranked according to essentiality score (bayes factor, log2), and WEE1 was identified as one of the top kinases in FUS::DDIT3-expressing SCP-1 cells, compared to YAP1 as established reference liability of MLS cells (33); FDR: False discovery rate. **C** ssGSEA pathway analysis of SCP-1 EV (green) and SCP-1 FUS::DDIT3 (blue) screening hits. Ten most essential pathways in FUS::DDIT3-expressing cells ranked according to normalized enrichment score (NES) are highlighted; ns: not significant.

To translate the results of the shRNA screen in genetically engineered SCP-1 cells into the setting of endogenous FUS::DDIT3 fusion protein expression in MLS, we assessed the protein levels and activity status of relevant WEE1 pathway components. Immunoblot analyses demonstrated increased WEE1 activity, indicated by enhanced WEE1-mediated inhibitory phosphorylation (P^Tyr15^) on CDK1 in FUS::DDIT3-expressing SCP-1 and representative MLS cells (Figure 2A). This was further accompanied by enhanced activity of the WEE1 upstream effectors CHK1 and CHK2, as indicated by increased P^Ser345^-CHK1 phosphorylation levels, implying an active replication stress response, and elevated P^Thr68^-CHK2 phosphorylation, demonstrating active ATM-CHK2 signaling. In line, we detected enhanced phosphorylation levels of RPA32 (P^Ser8^-RPA32; P^Ser33^-RPA32), serving as marker of replication stress. Combined with the detected elevated levels of γH2AX (P^Ser139^-H2AX), typically observed in the context of DNA damage, this indicated an elevated cellular stress level in FUS::DDIT3-expressing cells. Together, these results indicate that FUS::DDIT3-expressing cells are dependent on WEE1 activity.

**Figure 2.**
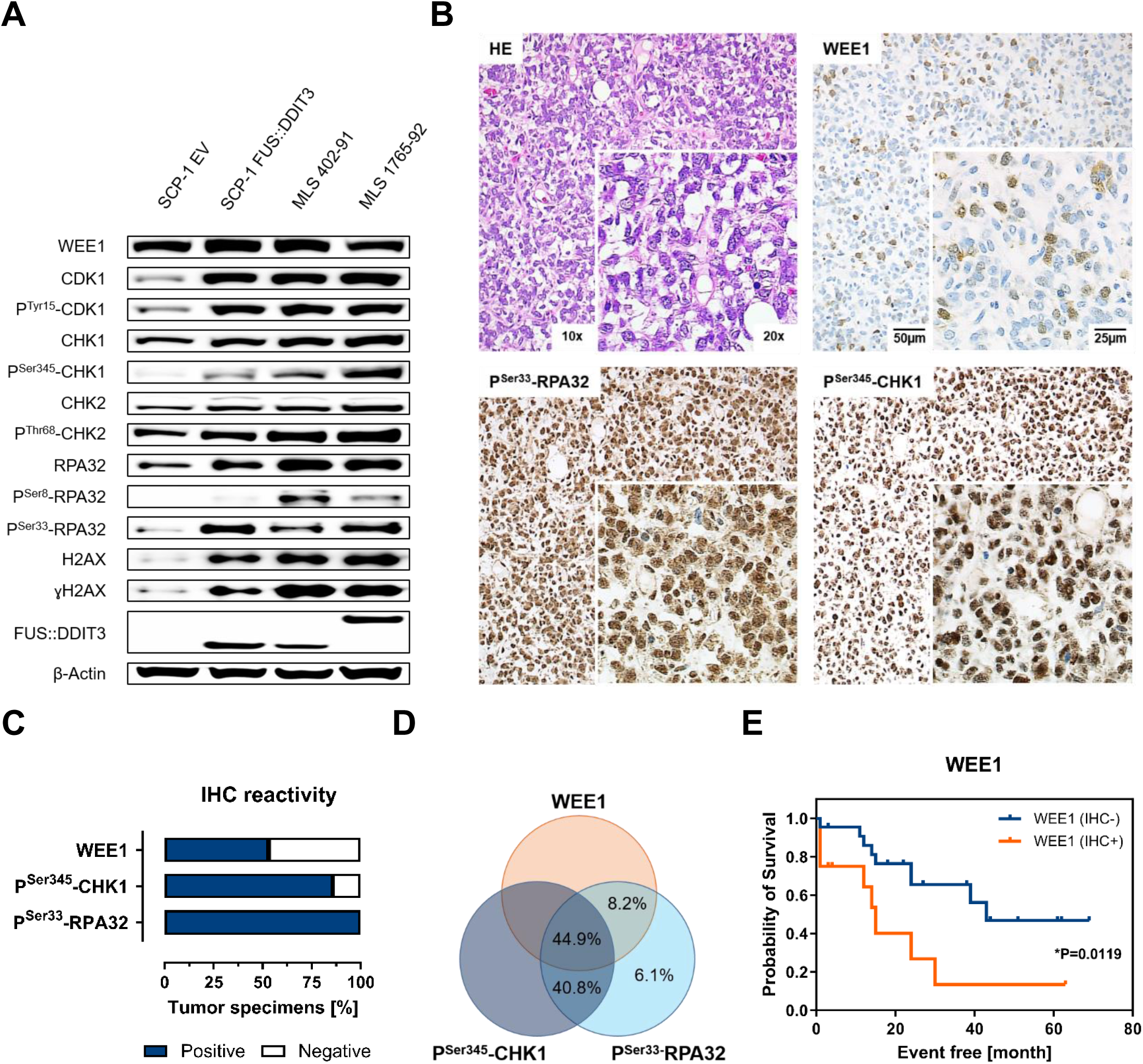
Enhanced WEE1 expression and pathway activity in FUS::DDIT3-expressing cells and MLS patient samples. **A** Immunoblot analysis of WEE1 signaling pathway activity in FUS::DDIT3-expressing and EV-transduced SCP-1 cells compared to representative MLS cell lines. One of at least two independent experiments with similar results is shown. **B** Representative immunoreactivity indicating strong expression of WEE1, P^Ser345^-CHK1 (WEE1 upstream effector kinase) and P^Ser33^-RPA32 (replication stress marker) in MLS tumor tissue specimens (*n*=49; original magnification, ×10 [inset, ×20]). **C** Overall IHC reactivity for WEE1, P^Ser345^-CHK1 and P^Ser33^-RPA32 in MLS patient samples. **D** Venn diagram showing overall concordance of WEE1, P^Ser345^-CHK1 and P^Ser33^-RPA32 immunoreactivity in MLS tumor tissue specimens (*n*=49). **E** Kaplan-Meier correlation displaying reduced overall/event free survival probability of WEE1 IHC positive (IHC+) MLS cases; P: *P*-value.

### WEE1 expression and pathway activity in MLS patient samples

To explore the involvement of WEE1 in MLS pathogenesis, we examined the expression levels of WEE1 in a cohort of 49 primary human MLS tissue specimens. In accordance with the immunoblot results in FUS::DDIT3-expressing cell lines, representative immunohistochemical (IHC) analyses showed moderate to strong expression levels of WEE1 pathway components (Figure 2B). WEE1 immunoreactivity was detected in 53.1% (26/49) of cases, which was in line with strong immunoreactivity of P^Ser345^-CHK1 in 85.7% (42/49) of tumors (Figure 2C). Furthermore, all MLS tissue specimens showed strong P^Ser33^-RPA32 immunoreactivity, indicating enhanced levels of replication stress in MLS tumors. Expression of WEE1, P^Ser345^-CHK1 and P^Ser33^-RPA32 as detected by immunoreactivity was concordant in 44.9% of MLS patients (Figure 2D).

Statistical analysis comparing WEE1 immunoreactivity with individual clinicopathological parameters of MLS patients revealed that WEE1 IHC positivity is significantly correlated with the aggressive round cell (high grade) histology of MLS, χ²(1) = 16.52, P = 0.0001 (Supplementary Figure S1A and Supplementary Table S1-3). Round cell MLS demonstrated significantly reduced overall/event free survival probability compared to myxoid (low grade) MLS (Supplementary Figure S1B). Similarly, WEE1 IHC reactivity was significantly correlated with reduced probability of event free patient survival (Figure 2E), indicating a prognostic impact of WEE1 expression in MLS. Individual clinicopathological parameters and scorings are listed in Supplementary Table S1 and S2. These results additionally support the concept that the cell cycle regulator WEE1 represents an essential effector in FUS::DDIT3-expressing MLS cells with enhanced WEE1 activity potentially required to tolerate high levels of replication stress.

### WEE1 activity is required for genomic integrity and MLS cell survival

To verify previous screening results and further elucidate whether enhanced WEE1 signaling activity confers sensitivity towards WEE1 functional loss in MLS cells, we performed RNAi-mediated *WEE1* depletion experiments in two MLS cell lines. Upon silencing of WEE1 in MLS 402-91 and MLS 1765-92 cells, we observed a reduction of P^Tyr15^-CDK1 phosphorylation levels, indicating decreased WEE1 activity (Figure 3A). This was accompanied by elevated cellular stress (indicated by increased P^Ser8^-RPA32 and γH2AX levels) and an activation of the replication stress response pathway, demonstrated by enhanced CHK1 phosphorylation in WEE1 knockdown samples. Furthermore, we observed that RNAi-mediated loss of *WEE1* is significantly associated with reduced MLS cell viability (Figure 3B). Together, these results indicate that FUS::DDIT3-expressing MLS cells are dependent on WEE1 activity.

**Figure 3.**
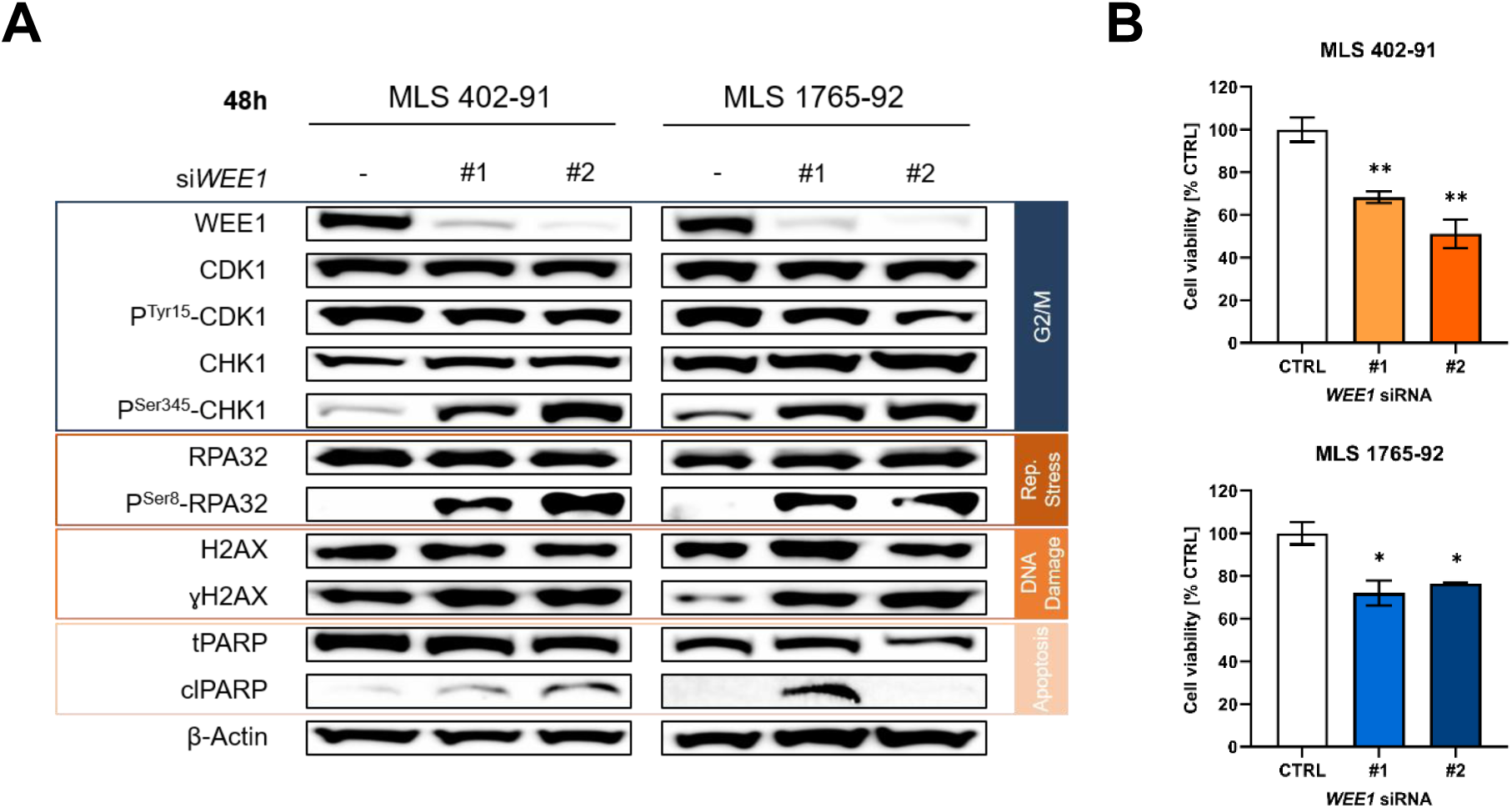
WEE1 activity is required for genomic integrity and MLS cell survival. **A** Expression analysis of P^Ser8^-RPA32 (replication stress markers) and ɣH2AX (DNA damage marker) following RNAi-mediated *WEE1* knockdown. One of at least two independent experiments with similar results is shown. **B** Cell viability assays of MLS 402-91 and MLS 1765-92 cells after RNAi-mediated *WEE1* knockdown. Bars and error bars represent the mean ± SEM of three independent experiments, *t*-test; CTRL: non-targeting control siRNA.

### MLS cells demonstrate sensitivity to pharmacologic inhibition of WEE1 activity

To evaluate the therapeutic potential of WEE1 inhibition in MLS, we first performed *in vitro* cell viability assays upon treatment with increasing concentrations of MK-1775, a small molecule ATP-competitive WEE1 inhibitor, for 72 h in various MLS cell lines in comparison to other liposarcoma subtypes. These experiments demonstrated dose-dependent suppression of cell viability in all cell lines (Figure 4A). A comparison of the calculated mean IC_50_ values of all analyzed cell lines, ranging from 0.19 µM to 2.88 µM (Supplementary Table S4), revealed that FUS::DDIT3-expressing MLS cell lines (highlighted in blue) are more sensitive to pharmacologic WEE1 inhibition compared to other liposarcoma subtypes. Associated dose-dependent reduction of WEE1 activity was confirmed in immunoblot analyses of MLS 402-91 cells, treated with increasing MK-1775 concentrations, through detection of decreased P^Tyr15^-CDK1 levels (Supplementary Figure S2A). MK-1775-mediated reduction of tyrosine 15 phosphorylation on CDK1 was further accompanied by activation of the CHK1 and CHK2 axes, as indicated by enhanced phosphorylation levels of both kinases (Figure 4B). This caused alternative G1/S cell cycle checkpoint activation via the p53-p21 axis in MLS cells, a potential cellular bypass mechanism. MK-1775 inhibitory effects were shown to be more prominent in the nuclear compartment of MLS protein fractions (Supplementary Figure S2B).

**Figure 4.**
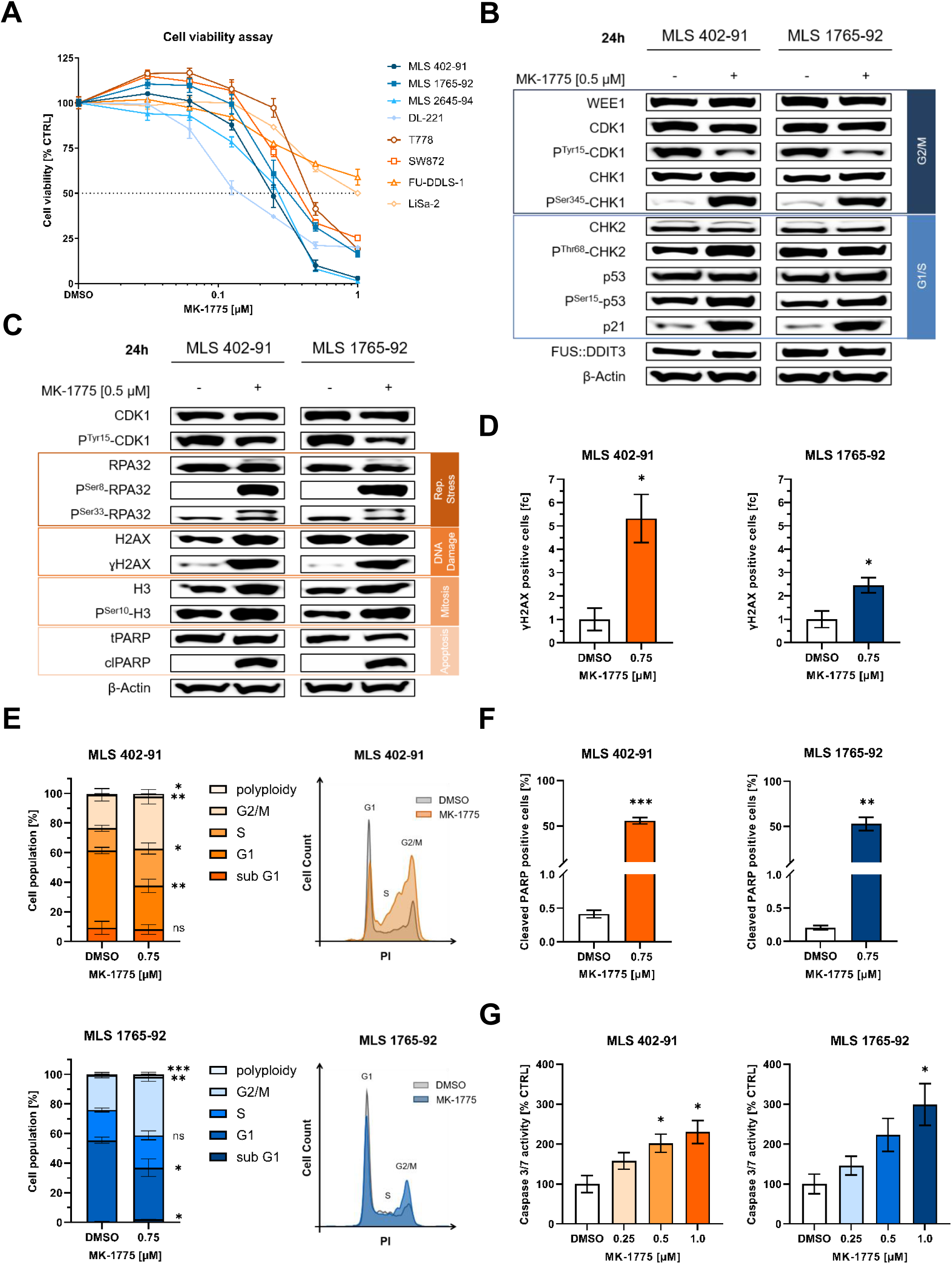
Sensitivity of MLS cells to pharmacologic inhibition of WEE1 activity. **A** Cell viability assays of liposarcoma cell lines (MLS: MLS 402-91, MLS 1765-92, MLS 2645-94, DL-221; ALT/WDLPS: T778; LPS: SW872; DDLPS: FU-DDLS-1; PLPS: LiSa-2) treated with MK-1775 for 72 h. Data points and error bars represent the mean ± SEM of one representative of three independent experiments measured in quintuplicate. **B** Immunoblot analysis of G2/M and G1/S-associated effectors in two MLS cell lines treated with 0.5 µM MK-1775 or DMSO (vehicle control) for 24 h. One of at least three independent experiments with similar results is shown. **C** Immunoblot analysis of replication stress (P^Ser8^-RPA32, P^Ser33^-RPA32), DNA damage (ɣH2AX), mitosis (P^Ser10^-H3) and apoptosis (cleaved PARP) in two MLS cell lines treated with 0.5 µM MK-1775 or DMSO (vehicle control) for 24 h. One of at least three independent experiments with similar results is shown. **D** Flow cytometry analysis of DNA damage (ɣH2AX) in two MLS cell lines treated with 0.75 µM MK-1775 or DMSO (vehicle control) for 24 h. Bars and error bars represent mean ± SEM of three independent experiments, *t*-test. **E** Flow cytometric analysis and representative cell cycle histograms of two MLS cell lines treated with 0.75 µM MK-1775 or DMSO (vehicle control) for 24 h. Bars and error bars represent the mean ± SEM of four independent experiments, *t*-test; ns: not significant; PI: Propidium iodide. **F** Flow cytometry analysis of apoptosis (cleaved PARP) in two MLS cell lines treated with 0.75 µM MK-1775 or DMSO (vehicle control) for 24 h. Values are presented as mean ± SEM of three independent experiments, *t*-test. **G** Caspase 3/7 activity assay of two MLS cell lines treated with increasing concentrations (0.25 – 1 µM) of MK-1775 or DMSO (vehicle control) for 24 h. Values are presented as mean ± SEM of three independent experiments, *t*-test.

Further immunoblot analysis demonstrated MK-1775-driven induction of cellular stress and DNA damage, as documented by enhanced RPA32 (P^Ser8^-RPA32; P^Ser33^-RPA32) phosphorylation and elevated γH2AX levels (Figure 4C). The DNA-damaged cell fraction was significantly elevated in MLS 402-91 cells and MLS 1765-92 cells upon MK-1775 treatment, as assessed by flow cytometric quantification (Figure 4D).

Despite previously detected G1/S cell cycle checkpoint activation and a high cellular stress level, MLS cells displayed mitotic capacity after pharmacologic WEE1 inhibition, as assessed by Ser10 phosphorylation levels of histone H3, a marker of mitosis, in immunoblots (Figure 4C). This is in line with results from flow cytometric cell cycle analysis of MLS cells demonstrating a significant drop in G1 cell fraction accompanied by an increase of cells entering S phase and a significant accumulation of cells in G2/M phase upon MK-1775 treatment, associated with cell cycle progression (Figure 4E). The significant increase in the G2/M fraction observed for MLS 402-91 and MLS 1765-92 cells either indicates mitotic arrest or premature mitotic entry.

As unresolved DNA damage accumulation can lead to induction of the apoptotic program and increased cell death, we further evaluated the apoptotic capacity of MK-1775 treatment by (i) analysis of PARP cleavage via immunoblotting as well as (ii) flow cytometric quantification and (iii) measurement of caspase 3/7 activity in luminescence-based apoptosis assays (Figure 4C, F, and G). Finally, as shown in Figures 4F and G, MLS cells revealed a dose-dependent induction of apoptosis upon MK-1775 treatment, as displayed by a significant increase of caspase 3/7 activity and elevated PARP cleavage compared to vehicle-control treated cells.

Collectively, our data reveal that genetic or pharmacologic inhibition of WEE1 activity causes enhanced replication stress and forces DNA-damaged MLS cells into premature mitosis ultimately leading to cell death via induction of apoptosis. Consequently, these results provide evidence that MLS cells depend on WEE1 to maintain cellular integrity and survival and that this reliance on WEE1 signaling represents a novel therapeutically addressable vulnerability of MLS.

### *In vivo* efficacy WEE1 inhibition in MLS CAM xenografts

To translate our findings and to confirm MK-1775 inhibitory efficacy *in vivo*, established MLS 402-91 and MLS 1765-92 CAM xenografts were treated with MK-1775 or appropriate vehicle control following evaluation of tumor growth reducing capacity (Figure 5). Topical administration of MK-1775 to MLS 402-91 xenografts resulted in a significant reduction of tumor volume with mean tumor regression of 45.8% compared to the vehicle-treated control group. Similar anti-proliferative effects were detected in MLS 1765-92 xenografts. Taken together, our *in vivo* data demonstrate that pharmacologic WEE1 inhibition impairs tumor growth, further substantiating the therapeutic potential of WEE1 inhibition in patients with FUS::DDIT3-driven MLS.

**Figure 5.**
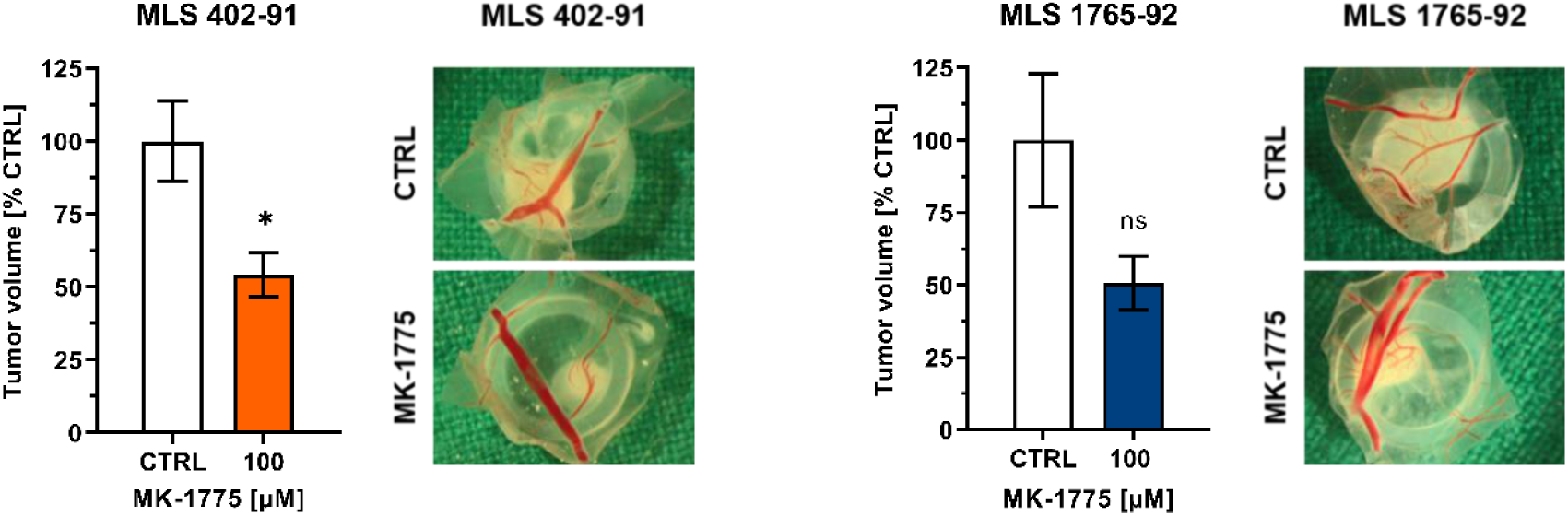
*In vivo* efficacy of WEE1 inhibition in MLS CAM xenografts. Tumor growth of MLS cells lines on chicken embryo CAMs following treatment with MK-1775 or DMSO. Shown are tumor volumes and representative images of tumors. Bars and error bars represent the mean ± SEM of at least three tumors, *t*-test; ns: not significant.

### FUS::DDIT3 fusion oncoprotein confers sensitivity to pharmacologic WEE1 inhibition

Since other fusion oncogene-expressing tumor entities reportedly depend on WEE1 pathway activity to counteract fusion oncoprotein-induced replication stress (34–36), we next evaluated whether the pathognomonic FUS::DDIT3 fusion protein contributes to the observed WEE1 inhibitor sensitivity in MLS. In proliferation assays performed in SCP-1 cells either expressing the FUS::DDIT3 fusion protein or an empty control vector treatment with MK-1775 resulted in significantly decreased cell viability. However, at lower MK-1775 doses this was exclusively observed in FUS::DDIT3-expressing cells (Figure 6A). The observed sensitivity of SCP-1 FUS::DDIT3 cells corresponds to high levels of replication stress, as indicated by P^Ser8^-RPA32, enhanced DNA damage, as displayed by γH2AX, and strong replication stress response, as indicated by P^Ser345^-CHK1, each detected upon MK-1775 treatment (Figure 6B). Furthermore, immunoblotting demonstrated amplified G1/S cell cycle checkpoint signaling, as indicated by enhanced P^Thr68^-CHK2 and P^Ser15^-p53 levels, particularly prominent in SCP-1 FUS::DDIT3 cells. To evaluate potential differences in cell cycle checkpoint regulation between SCP-1 EV and FUS::DDIT3 cells, flow cytometric cell cycle analysis was performed in MK-1775-treated SCP-1 cells and an appropriate control. Results suggested that expression of FUS::DDIT3 alone drives S phase entry, indicated by stronger reduction in G1 cell fraction and concurrent 3-fold enhanced S phase fraction in FUS::DDIT3-expressing cells compared to SCP-1 EV cells, determined in the control setting. Interestingly, FUS::DDIT3-expressing cells maintained stronger cell cycle progression upon MK-1775 treatment, indicated by a more prominent reduction of the G1 cell fraction, accompanied by elevated S and G2/M cell fractions, whereas SCP-1 EV cells demonstrated a minor decrease in the G1 phase fraction and primarily accumulated in S phase (Figure 6C). Correspondingly, apoptosis assays measuring caspase 3/7 activity showed that lower MK-1775 concentrations are sufficient to significantly induce apoptosis in FUS::DDIT3-expressing SCP-1 cells, whereas similar inhibitor concentrations demonstrated minor effects in the control cell line (Figure 6D).

**Figure 6.**
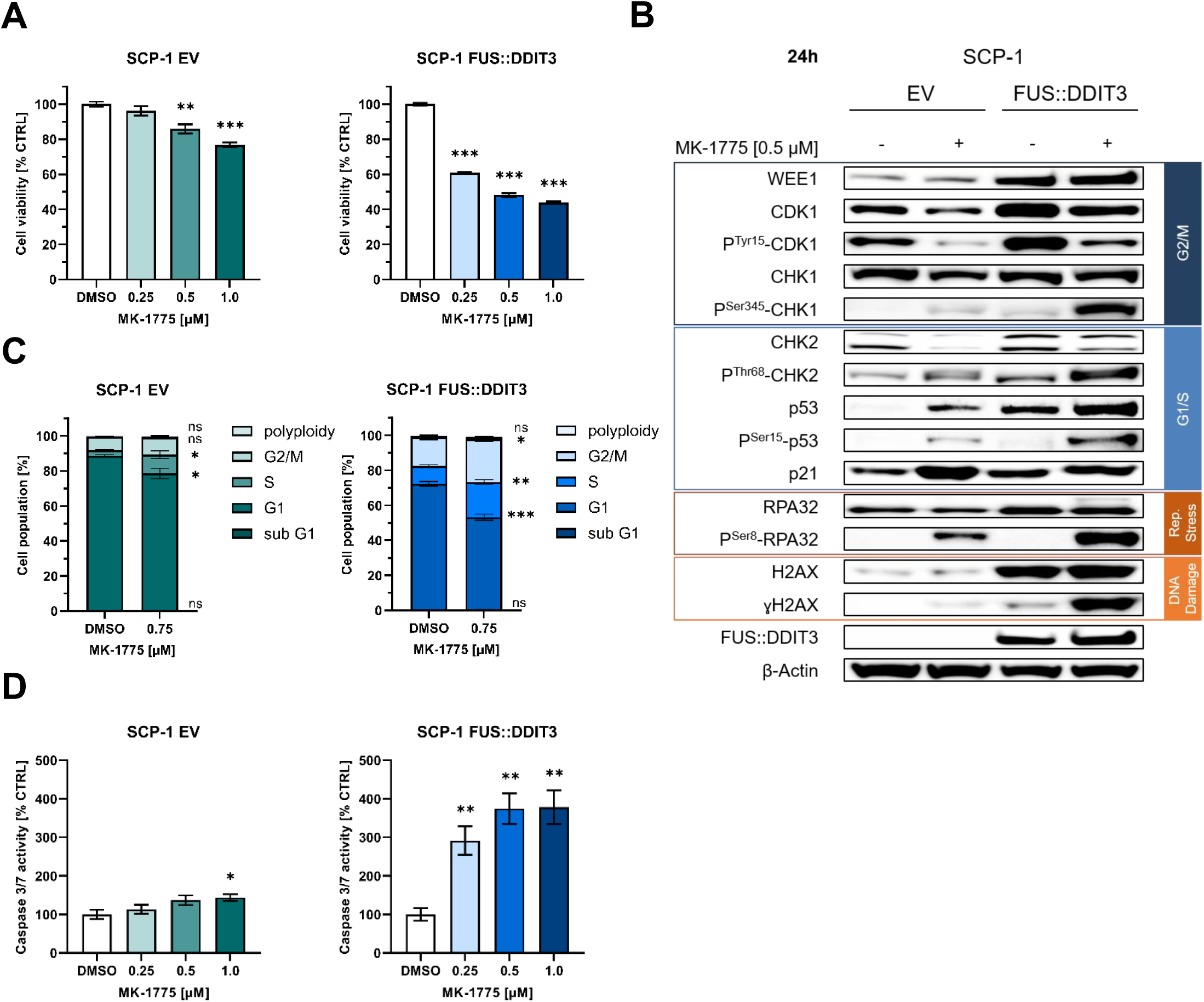
The FUS::DDIT3 fusion oncoprotein confers sensitivity to pharmacologic inhibition of WEE1 activity. **A** MTT cell viability assays of FUS::DDIT3-expressing or EV-transduced SCP-1 cells cultured in the presence of increasing MK-1775 concentrations (0.25 – 1 µM) for 72 h. DMSO was applied as vehicle control. Data points and error bars represent the mean ± SEM of three independent experiments, *t*-test. **B** Immunoblot analysis of cell cycle checkpoints, replication stress (P^Ser8^-RPA32) and DNA damage (ɣH2AX) in SCP-1 EV and FUS::DDIT3 cells treated with 0.5 µM MK-1775 or DMSO (vehicle control) for 24 h. One of at least three independent experiments with similar results is shown. **C** Cell cycle distribution assessed by flow cytometric propidium iodide staining of SCP-1 cells treated with MK-1775 (0.75 µM) or DMSO for 24 h. Data are presented as mean ± SEM of three independent experiments, *t*-test; ns: not significant. **D** Caspase 3/7 activity assay in SCP-1 EV and FUS::DDIT3 cells treated with increasing concentrations (0.25 – 1 µM) of MK-1775 or DMSO (vehicle control) for 24 h. Values are presented as mean ± SEM of three independent experiments, *t*-test.

Collectively, our data provide first evidence that sensitivity towards loss of WEE1 activity in MLS is mediated by the chimeric FUS::DDIT3 fusion oncoprotein, potentially via enhanced FUS::DDIT3-driven G1/S cell cycle transition.

### Causal relationship between FUS::DDIT3-modulated G1/S cell cycle checkpoint regulation and requirement for WEE1 activity

Our observations indicate that FUS::DDIT3 drives S phase entry potentially via G1/S cell cycle checkpoint deregulation. This reportedly causes accelerated S phase entry and enhanced DNA replication, which promotes replication stress, G1/S deregulation confers functional G2/M checkpoint dependence in other tumor cells (37). Therefore, we investigated the contribution of Cyclin E and its kinase CDK2 as key mediators of G1/S cell cycle transition to observed WEE1 requirement in MLS cells. We first assessed the general expression level of Cyclin E1 and E2 and the activity status of CDK2 in SCP-1 and MLS cells via immunoblotting. Results demonstrated FUS::DDIT3-dependent expression of Cyclin E1 and enhanced CDK2 activity, indicated by elevated P^Thr160^-CDK2 levels, in SCP-1 as well as MLS cells (Figure 7A). Examining expression levels of Cyclin E1 and P^Thr160^-CDK2 in a cohort of MLS tissue specimens (n=49) via IHC we observed Cyclin E1 immunoreactivity in 63.3% (31/49) of MLS specimens, and 67.3% (33/49) of samples revealed P^Thr160^-CDK2 expression (Figure 7B), consistent with previous immunoblot results. Concurrent expression of both markers was detected in 46.9% (23/49) of MLS specimens and a prominent overlap in immunoreactivity of the WEE1 axis and G1/S regulators was observed in 26.5% (13/49) of MLS tumors (Supplementary Figure S3A). Statistical analysis comparing immunoreactivity with individual clinicopathological parameters of MLS patients revealed that P^Thr160^-CDK2 IHC positivity is significantly correlated with (i) positive WEE1 immunoreactivity, χ²(1) = 11.23, P = 0.002 (Supplementary Table S3), (ii) the more aggressive round cell histology of MLS (Supplementary Figure S3B) and (iii) reduced overall/event free survival in Kaplan-Meier correlations (Figure 7C).

**Figure 7.**
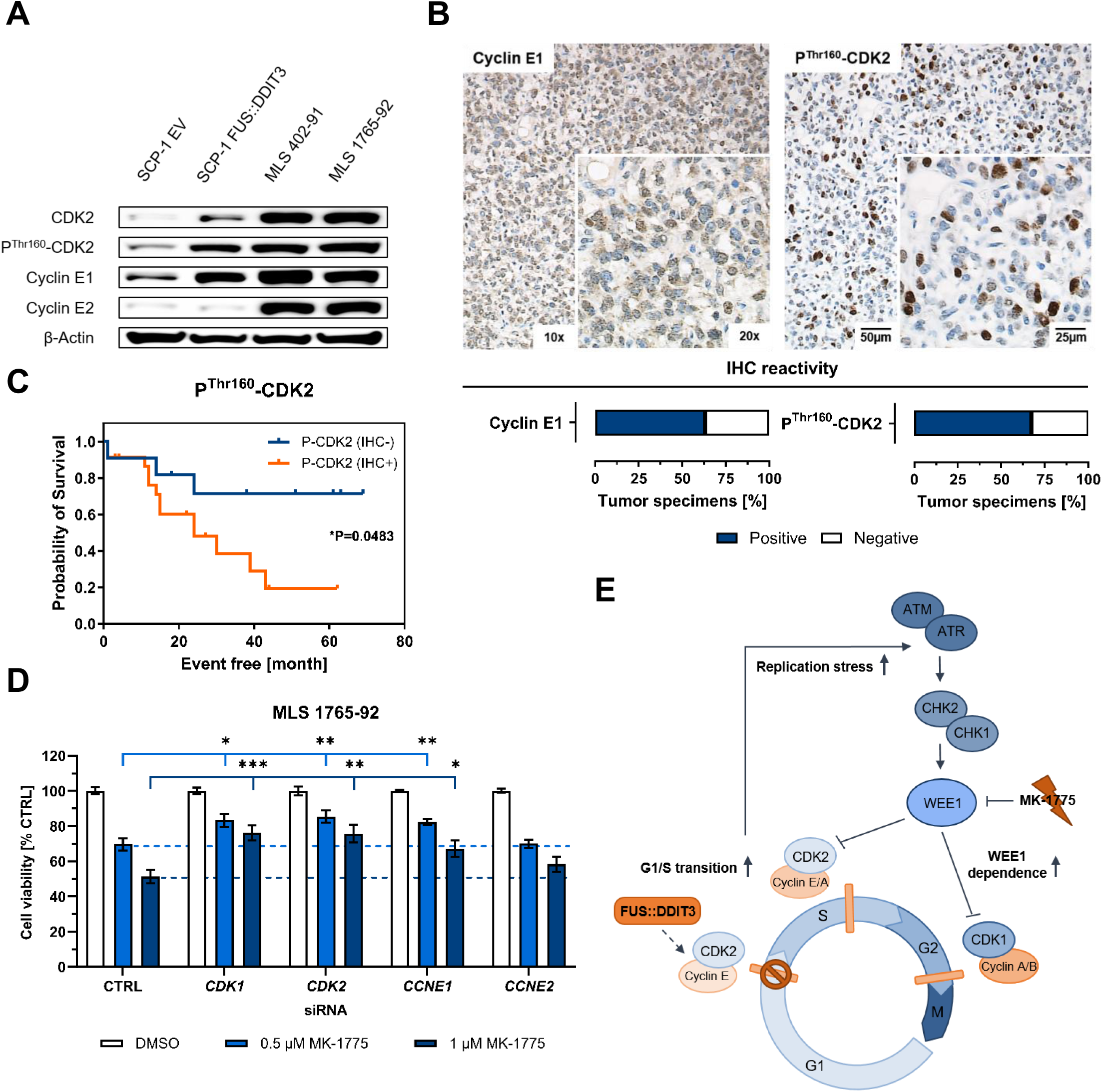
Causal relationship between FUS::DDIT3-modulated G1/S cell cycle checkpoint regulation and requirement for WEE1 activity. **A** Immunoblot analysis of CDK2 and Cyclin E1/2 expression in SCP-1 and MLS cells. **B** Representative immunostains documenting active CDK2 (P^Thr160^-CDK2) and Cyclin E1 in MLS tumor tissue specimens (n=49; original magnification, ×10 [inset, ×20]). **C** Kaplan-Meier correlation showing reduced overall/event free survival probability of P^Thr160^-CDK2 IHC positive (IHC+) MLS cases; P: *P*-value. **D** Cell viability assay of MLS 1765-92 cells after RNAi-mediated depletion of *CDK1*, *CDK2*, *CCNE1* or *CCNE2* and additional treatment with MK-1775 (0.5 – 1 µM) or DMSO (vehicle control). One representative of three independent experiments presented as mean ± SEM is shown. **E** Schematic overview of suggested FUS::DDIT3-driven G1/S cell cycle checkpoint deregulation via Cyclin E1/CDK2 overexpression causing cell cycle transition into S phase, replication stress and enhanced requirement for WEE1 pathway to maintain intra-S and G2/M cell cycle checkpoint activity in MLS cells.

We further explored whether G1/S cell cycle checkpoint deregulation via highly active Cyclin E/CDK2 represents the decisive factor for WEE1 inhibitor sensitivity in FUS::DDIT3-expressing cells. Therefore, a *CDK* and *CCNE* siRNA-mediated depletion approach was performed in MLS 1765-92 cells, followed by 24 h treatment with increasing MK-1775 concentrations or vehicle control. Effects were assessed via cell proliferation analysis and immunoblotting. MTT assays demonstrated that RNAi-mediated *CDK1* and *CDK2* depletion, as well as decreased *CCNE1* expression levels were sufficient to rescue cell viability upon MK-1775 treatment in MLS 1765-92 cells (Figure 7D). However, *CCNE2* depletion had a minor effect on MK-1775 sensitivity. Immunoblotting further showed that *CDK2* knockdown alone completely abrogates MK-1775-induced replication stress, indicated by complete loss of RPA32 phosphorylation and substantial decrease of Ser345 phosphorylation of replication stress response marker CHK1 (Supplementary Figure S4). In addition, *CDK2* silencing dampens DNA damage and apoptosis, displayed by γH2AX and cleaved PARP, respectively. For *CCNE1* depleted MLS cells, mitigated effects with a similar tendency were observed.

Collectively, our results suggest a model in which FUS::DDIT3-driven expression of Cyclin E1 and its kinase CDK2 impairs G1/S cell cycle checkpoint function and thereby confers therapeutic dependence on G2/M cell cycle checkpoint and WEE1 activity in MLS cells (Figure 7E).

The present study uncovered WEE1 kinase activity as functional vulnerability of FUS::DDIT3-expressing MLS cells and provides first evidence that interference with WEE1 signaling represents a novel promising strategy for therapeutic intervention in MLS.

## Discussion

MLS represent a group of rare malignant neoplasms that display a high rate of local recurrence and metastasis (6). While localized disease is associated with improved prognosis with up to 85% estimated 10-year survival rates owing to a large extent to enhanced radio- and chemotherapeutic responsiveness, prognosis declines rapidly in the advanced setting (38–40), underlining the urgent need for novel therapeutic approaches – especially in the management of advanced and metastatic disease.

Since MLS cells harbor only few additional mutations aside from the pathognomonic fusion protein (41, 42), which is, so far, not pharmacologically addressable, the most promising strategy to therapeutically target MLS represents interference with FUS::DDIT3-driven oncogenic effectors and signaling pathways. Based on a functional genomic screen (33), the cell cycle checkpoint kinase WEE1, an important component of the highly conserved replication stress response machinery, was identified as FUS::DDIT3-essential effector Here we investigated the relevance of WEE1 in FUS::DDIT3-expressing MLS cells and the underlying mechanism for enhanced requirement for WEE1 pathway activity.

Oncogenic fusion genes encoding aberrant transcription factors are drivers of replication stress (43, 37), as shown in previous studies in PAX::FOXO1-driven rhabdomyosarcoma, CIC::DUX4-expressing sarcoma and Ewing sarcoma harboring EWSR1::FLI1 fusion oncoproteins (34–36). Consistently, FUS::DDIT3 expression in a mesenchymal stem cell model system was sufficient to accelerate cell cycle progression and to induce cellular stress. Enhanced cell proliferation and associated replication stress requires functional cell cycle checkpoint signaling to prevent propagation of genetic errors and DNA damage accumulation (44–46). This aligns with results from the present pathway analysis showing that FUS::DDIT3-expressing cells functionally depend on genes involved in the cell cycle machinery, DNA replication and mitosis. Increased WEE1 and replication stress response pathway activity, noted in MLS cell lines and tissue specimens, was hence suggested to represent a pro-oncogenic mechanism to tolerate induced proliferative drive while preventing lethal levels of replication stress, maintaining genomic integrity and cellular survival. This corresponds to our observation that WEE1 expression in MLS tumor specimens is significantly correlated with the more aggressive round cell histology of MLS and equally contributes to reduced overall/event free survival in our MLS patient cohort. In line with the present findings, WEE1 expression was significantly associated with worse disease-free survival in melanoma (27), poor overall survival in ovarian carcinoma (47) and higher risk grade in GIST (48). This suggests an involvement of WEE1 kinase activity in MLS progression and implies its potential as a prognostic marker, which needs to be further explored in prospective studies.

According to the indicated functional relevance of WEE1 activity in MLS pathogenesis, we demonstrate, either by genetic silencing or pharmacologic inhibition with the small molecule inhibitor MK-1775, that *in vitro* and *in vivo* models of MLS are highly sensitive to loss of required WEE1 kinase signaling. Previous results in other sarcoma entities demonstrated that MK-1775-forced premature mitotic entry of cells with accumulated DNA damage ultimately causes cell death via mitotic catastrophe (49). Correspondingly, our data revealed, that in FUS::DDIT3-expressing cells pharmacologic inhibition and RNAi-mediated genomic depletion induce G2/M cell cycle checkpoint abrogation, enabling cell cycle progression in the presence of enhanced replication stress and DNA damage which resulted in an induction of the apoptotic program. Strong discrepancies between MK-1775-mediated effects on cell viability, replication stress and DNA damage, as well as apoptosis in SCP-1 cells, either expressing the fusion protein or the control vector, confirmed that FUS::DDIT3 fusion oncoprotein activity sensitizes cells to WEE1 inhibition. Hence, our study provides evidence that WEE1 represents a promising FUS::DDIT3-dependent therapeutic vulnerability in MLS.

Aiming at an elucidation of the potential underlying FUS::DDIT3-driven mechanisms that contribute to the observed WEE1 requirement, we found prominent differences in G1/S cell cycle transition between FUS::DDIT3- and EV-expressing SCP-1 cells. Interestingly, deregulatory mechanisms in cell cycle modulators affecting G1/S transition are commonly observed in liposarcoma. While WDLPS and DDLPS comprise *CDK4* and *MDM2* amplifications, PLPS are characterized by a more complex mutational status including loss of *RB1* or *TP53* mutations contributing to a defective G1/S cell cycle checkpoint and dependence on functional G2/M signaling (50, 51, 41, 52). Conversely, in MLS similar mutations arise only sporadically (53) and observed *TP53* mutations in DL-221 cells represent a unique feature among the MLS cell lines included in the present study (54). Despite this, all FUS::DDIT3-expressing MLS cells demonstrated higher responsiveness to MK-1775 compared to other types of liposarcoma in MTT assays. Therefore, an alternative FUS::DDIT3-driven G1/S deregulation mechanism promoting WEE1 pathway requirement in MLS was assumed. In line with published studies indicating that alternative G1/S cell cycle checkpoint deregulation mechanisms and factors are more suitable to predict responsiveness (55, 56), we focused on Cyclin E and its kinase CDK2 as key mediators of G1/S cell cycle transition, since an aberrant activation of the Cyclin E/CDK2 complex contributes to increased replication origin firing and DNA replication defects causing high levels of replication stress and genomic instability (57, 58). This also coincides with the observed FUS::DDIT3-mediated replication stress phenotype in MLS.

In line with previous reports of aberrant expression of G1 controlling cyclins, Cyclin E1, Cyclin D1 and their kinases CDK2 and CDK4 observed in MLS tissues (59), we showed enhanced FUS::DDIT3-dependent expression of Cyclin E1 and concurrent CDK2 activity in SCP-1 and MLS cells, as well as MLS tumor specimens in the present study. Providing a potential explanation, recently published results by Kadoch et al. revealed FUS::DDIT3-driven expression of gene sets implicated in cell cycle, G2/M checkpoint and E2F pathways in MLS cells, that were similar to expression patterns of primary MLS tumors (60). Additional findings of our group in MLS cells demonstrated FUS::DDIT3-driven transcriptional upregulation of E2F target genes, including Cyclin E1, and thus provide further evidence for a FUS::DDIT3-induced dysregulation of the G1/S cell cycle checkpoint via the E2F axis (61). However, a direct fusion protein-mediated transcriptional upregulation of target genes, as reported for CIC::DUX4-driven sarcomas, demonstrating fusion oncoprotein-mediated Cyclin E1 expression due to occupation of *CCNE1* promoter regions, is also conceivable (62). In MLS, a similar mode of action has already been described for the transcription of *IGF2*, which was shown to be mediated via (direct or indirect) action of FUS::DDIT3 on the *IGF2* P2 promoter (63).

Corresponding to their functional implication in replication stress, aberrant Cyclin E/CDK2 activity reportedly sensitizes tumor cells to pharmacologic WEE1 inhibition (64–67, 55). Such a causal relationship is consistent with the observed correlation between CDK2 activity and WEE1 immunoreactivity in our MLS patient cohort. Moreover, we were able to show that MK-1775 responsiveness in MLS is dependent on CDK1, CDK2 and Cyclin E1 expression, which aligns with published results from experiments in CIC-rearranged sarcomas (35). The prominent rescue of MLS cell viability upon MK-1775 treatment in both, *CDK1* and *CDK2*-depleted cells, can be explained by WEE1 dual specificity for the G2/M and intra-S cell cycle checkpoint. However, immunoblotting showed that MK-1775-induced replication stress, DNA damage and apoptosis were predominantly dependent on CDK2 and to a lesser extent on Cyclin E1 expression. This is in line with data from a similar study showing that Cyclin E1-mediated replication stress and DNA damage is dependent on functional CDK2 (64). These findings imply that substantial MK-1775 cytotoxic effects are promoted via the CDK2 axis in S phase in a G2/M-independent manner, a notion supported by other studies (68, 69, 32). Further highlighting a functional relevance of CDK2 in FUS::DDIT3-driven oncogenesis in MLS, findings by Bento et al. confirmed direct interaction of CDK2 with the FUS::DDIT3 fusion protein via its DDIT3 portion, potentially influencing the functions of both binding partners (70).

Our observations provide first evidence that IHC screening for Cyclin E and CDK2 might offer a novel opportunity to predict therapeutic MLS’ responsiveness to MK-1775 treatment requiring further investigation in prospective studies. In line, *CCNE1* amplification/overexpression was identified as biomarker in other tumor entities and already proved applicability in patient selection for MK-1775 clinical trials in refractory solid tumors (71, 67). Given the high complexity of cell cycle checkpoint regulation we are aware that several other biomarkers might be similarly useful in MK-1775 response prediction, as shown for homologous recombination repair defects (e.g. BRCA1/2 inactivation) in other soft tissue sarcoma subtypes (72).

In conclusion, we have identified the cell cycle checkpoint kinase WEE1 activity as FUS::DDIT3-dependent functional requirement in MLS cells. Our findings provide first evidence that requirement for active WEE1 signaling, maintaining a functional G2/M and intra-S cell cycle checkpoint, is mediated via FUS::DDIT3 fusion oncoprotein-driven upregulation of the cell cycle regulators Cyclin E1 and CDK2. Our data contribute to a better understanding of the FUS::DDIT3 fusion protein’s oncogenic effects driving MLS pathogenesis and provides first preclinical evidence that WEE1 activity represents a rational target as basis for biology-guided therapeutic intervention in MLS.

## Material and Methods

### Cell culture

The human myxoid liposarcoma cell lines MLS 402-91 (*FUS::DDIT3* exon 7::2) MLS 2645-94 (*FUS::DDIT3* exon 5::2) and MLS 1765-92 (*FUS::DDIT3* exon 13::2) (contributed by Pierre Åman), as well as well-differentiated liposarcoma (WDLPS) cell line T778 (contributed by Florence Pedeutour), dedifferentiated liposarcoma (DDLPS) FU-DDLS-1 cells (contributed by Jun Nishio), pleomorphic liposarcoma (PLPS) cell line LiSa-2 (contributed by Marcus Renner) and liposarcoma (LPS) cell line SW872 (obtained from CLS Cell Lines Service) were cultured in RPMI 1640 medium supplemented with 10% fetal bovine serum (FBS; Gibco). DL-221 cells (*FUS::DDIT3* exon 7::2) were cultured in DMEM medium (10% FBS). The human mesenchymal SCP-1 stem cell system, established and contributed by Thomas Kindler (33), was cultured in MEM medium (10% FBS). All cell lines were cultured under standard conditions (37°C and 5% CO_2_ in a humidified atmosphere) for a maximum of 20 passages and frequently tested for mycoplasma contamination. Cell line identity was verified by cell authentication SNP profiling (Multiplexion) and/or validation of the specific *FUS::DDIT3* gene fusion via RT-PCR. To study the effect of small molecule inhibition (MK-1775), cell lines were grown in medium supplemented with 2% FBS.

### shRNA screen re-analysis and ssGSEA

Raw read counts were normalized, and log2 fold-change (FC) per shRNA was calculated using plasmid DNA as a baseline. Then, shRNA was selected to the three best shRNA per gene, and the log2-bayes factor (essentiality score) was calculated using CEG1 and non-essential gene set as reference. The re-analysis was performed using the BAGEL algorithm (73).

To perform gene set enrichment analysis (GSEA), we calculated the differential essentiality score by subtracting the essentiality score (FUS::DDIT3 - EV) and transforming it to z-score. The result was used as input for ssGSEA 2.0 R package.

### Tumor specimens and tissue microarrays

Tissue microarrays (TMA) were prepared from 49 formalin-fixed and paraffin-embedded MLS specimens selected from the archive of the Gerhard-Domagk-Institute of Pathology (Münster University Hospital, Münster, Germany). Differential diagnoses were confirmed by break-apart FISH or RNA sequencing and reviewed by two experienced pathologists based on current World Health Organization (WHO) criteria. The study was approved by the Ethics Review Board of the University of Münster (2015-548-f-S), and experiments were conformed to the principles set out in the World Medical Association Declaration of Helsinki and the United States Department of Health and Human Services Belmont Report.

### Immunohistochemistry

Immunohistochemistry (IHC) staining was performed with a Bench-Mark ULTRA Autostainer (VENTANA/Roche) on 3 µm TMA sections. The staining procedure included heat-induced epitope retrieval using Tris-Borate-EDTA buffer (pH 8.4; 95-100°C, 32-72 min) followed by primary antibody labelling (16-120 min) and signal detection carried out with the OptiView DAB IHC Detection Kit (VENTANA/Roche), as described previously (33). All applied primary antibodies are listed in Supplementary Table S5. Immunoreactivity was evaluated applying a staining intensity and a proportion score. Staining intensity was assessed by a semi-quantitative score (0, negative; 1, weak; 2, moderate; and 3, strong) defined by the strong staining intensity of the positive control tissue (colorectal cancer; score = 3). In order to assess the proportion score, the following categories were applied: 1 = 0-24%, 2 = 25-49%, 3 = 50-74% and 4 = 75-100%. The signal intensity score and proportion score were multiplied and the cut-off score, considering tumor specimens as positive for the purpose of the study, was set to ≥ 6. WEE1 immunoreactivity was considered positive, if ≥ 5% of cells demonstrated moderate signal intensity. The scoring system was pre-specified without prior analyses of the clinical course and the IHC readers were blinded to outcome data.

### RNA interference

Short interfering RNA (siRNA)-mediated silencing of *WEE1*, *CCNE1*, *CCNE2*, *CDK1* and *CDK2* was performed with a set of siRNAs purchased from Life Technologies: *WEE1* #1 (ID 406, pre-validated), *WEE1* #2 (ID 103582), *CCNE1* (ID s2526, pre-validated), *CCNE2* (ID s17450, pre-validated), *CDK1* (ID 1625, pre-validated) and *CDK2* (ID 1314, pre-validated). Block-iT Alexa Fluor Red Fluorescent Control siRNA was used as non-targeting negative control (Life Technologies, #14750100). Cells were grown in 60 mm cell culture dishes in medium supplemented with 2% FBS overnight and transfected with 62.5 pmol of the indicated siRNAs applying Lipofectamine RNAiMAX (Invitrogen, #13778150). After 48-72 h, cells were harvested and transfection efficiency was determined via immunoblotting.

### Compounds

The ATP-competitive WEE1 inhibitor MK-1775 (C_27_H_32_N_8_O_2_; CAS#: 955365-80-7) (29) was purchased from Selleckchem (#S1525) and dissolved in DMSO (AppliChem, A3672). The final DMSO concentration of the applied DMSO vehicle control did not exceed 0.2% (v/v) in all *in vitro* and *in vivo* experiments. Dosing of MK-1775 was aligned to a Phase I study in patients with advanced solid tumors (71).

### Cell lysate preparation and immunoblotting

Cells were harvested and lysed with 1x RIPA buffer (CST, #9806) supplemented with 1x protease inhibitor cocktail (Roche, #693116001) and 1x phosphatase inhibitor cocktail (Roche, #04906845001) according to the manufacturer’s instructions. Whole cell protein lysates were separated by SDS-PAGE and transferred to a nitrocellulose membrane via western blotting. Membranes were blocked with 5% dry milk or 5% BSA in TBST and incubated with primary antibodies listed in Supplementary Table S5. Targets were labelled with HRP-linked secondary antibodies and chemiluminescent signal detection was carried out with the Signalfire ECL Reagent and the Molecular Imager ChemiDoc System (Image Lab Software, version 5.1, Bio-Rad Laboratories).

### Flow cytometry analysis

For cell cycle analysis, MLS cells were incubated in medium supplemented with 2% FBS and indicated concentrations of MK-1775 or vehicle control for 24 h. In brief, cells were trypsinized, washed in PBS containing 0.5% BSA and permeabilized with ice-cold ethanol for 30 min. Subsequently, cells were treated with RNase A (Merck Millipore, #708563) and stained with propidium iodide (PI) solution (Sigma-Aldrich, #P4170) for 10 min at room temperature. For measurement of MK-1775-induced DNA damage or apoptosis, cells were fixed in 2% paraformaldehyde (PFA) for 10 min, permeabilized and stained with conjugated antibodies listed in Supplementary Table S5 at room temperature for 1 h. Cells were measured on a BD FACS Celesta cytometer and data analysis was performed with FlowJo software (version 10.8.1). At least three independent experiments were performed.

### Cell viability assay

To determine the effects of pharmacologic WEE1 inhibition on MLS cell viability, MLS 402-91 cells (2.5 x 10^3^), MLS 1765-92 cells (2.2 x 10^3^), MLS 2645-92 (2 x 10^3^), DL-221 (1.5 x 10^3^), T778 (2.5 x 10^3^), FU-DDLS-1 (5 x 10^3^), LiSa-2 (3.8 x 10^3^), SW872 (3.5 x 10^3^), SCP-1 EV (4 x 10^3^) and SCP-1 FUS::DDIT3 (4 x 10^3^) were seeded in 96-well cell culture plates (100 µL/ well; medium containing 2% FBS) and treated with increasing concentrations of MK-1775 (0.03125-4 µM) for 72 h. Cell viability was measured using the Cell Proliferation Kit I (MTT) (Roche, #11465007001) according to the manufacturer’s instructions. For RNAi-mediated knockdown experiments, cells were transfected with indicated siRNAs 72 h prior to MTT assays. At least three independent experiments were performed in quintuplicates.

### Apoptosis assay

The apoptotic effect of MK-1775 was determined by means of caspase 3/7 activity via Apotox-Glo Triplex Assay (Promega, #G6320) according to the manufacturer’s protocol. MLS cells (MLS 402-91: 1.5 x 10^4^; MLS 1765-92: 1.25 x 10^4^) and SCP-1 cells (EV: 1.25 x 10^4^; FUS::DDIT3: 1.6 x 10^4^) were seeded in 96-well cell culture plates and treated with increasing concentrations of MK-1775 (0.25-1 µM) for 24 h. At least three independent experiments were performed in quintuplicates.

### Chicken chorioallantoic membrane (CAM) assays

To evaluate MK-1775 *in vivo* efficacy, CAM assays were performed as previously described (74). In brief, MLS 402-91 and MLS 1765-92 cells (1.5 x 10^6^ cells/egg; dissolved in medium/ Matrigel 1:1, v/v) were deposited within 5 mm silicon rings on the surface of chorioallantoic membranes of fertilized chicken eggs and incubated at 37°C in a 60% humidified atmosphere. On days eight, nine and ten, MK-1775 or DMSO vehicle control (0.2% DMSO in 0.9% NaCl) was applied topically. Three days after treatment initiation, CAM xenografts were imaged, explanted, fixed in 5% PFA, and processed for histopathological examination. Tumor volume (mm^3^) was calculated according to the formula: length (mm) × width^2^ (mm) × π/6 (75).

### Statistical analysis

Computations were performed using GraphPad Prism software (version 9). Statistical difference was calculated using two-tailed Student’s *t*-test or Fisher’s exact test as appropriate and considered significant at *P*<0.05 (*), *P*<0.01 (**), and *P*<0.001 (***). Statistical correlation analysis was carried out by means of χ². Data points and error bars represent the mean ± SEM of *n* independently performed experiments unless otherwise stated.

## Acknowledgements

The authors thank Esther-Pia Jansen, Birgit Schulte, Inka Buchroth and Eva Winkler for excellent technical support. Thomas Kindler (University Cancer Center, University Medical Center of Mainz, Mainz, Germany) kindly provided the SCP-1 cells. Florence Pedeutour (Laboratory of Solid Tumor Genetics, Institute for Research on Cancer and Aging of Nice; Laboratory of Solid Tumor Genetics, University Hospital of Nice-Côte d’Azur University, Nice, France) kindly provided the T778 cell line. Marcus Renner (Division of Translational Medical Oncology, National Center for Tumor Diseases (NCT) Heidelberg and German Cancer Research Center (DKFZ), Heidelberg, Germany) kindly provided the LiSa-2 cell line. Jun Nishio (Faculty of Medicine, Department of Orthopaedic Surgery, Fukuoka University, Fukuoka, Japan) kindly provided the FU-DDLS-1 cell line. This study was supported in part by grants from the Deutsche Krebshilfe (M. Trautmann and S. Fröhling; 70113604 and 70113610), the “Innovative Medical Research” funding program of the University of Münster Medical School (M. Trautmann; TR122011) and the Deutsche Forschungsgemeinschaft / DFG (W. Hartmann and M. Trautmann, HA4441/2-1).

