## Supplementary Material for "Exploiting WEE1 kinase activity as FUS::DDIT3-dependent therapeutic vulnerability in myxoid liposarcoma"

### Slide 1
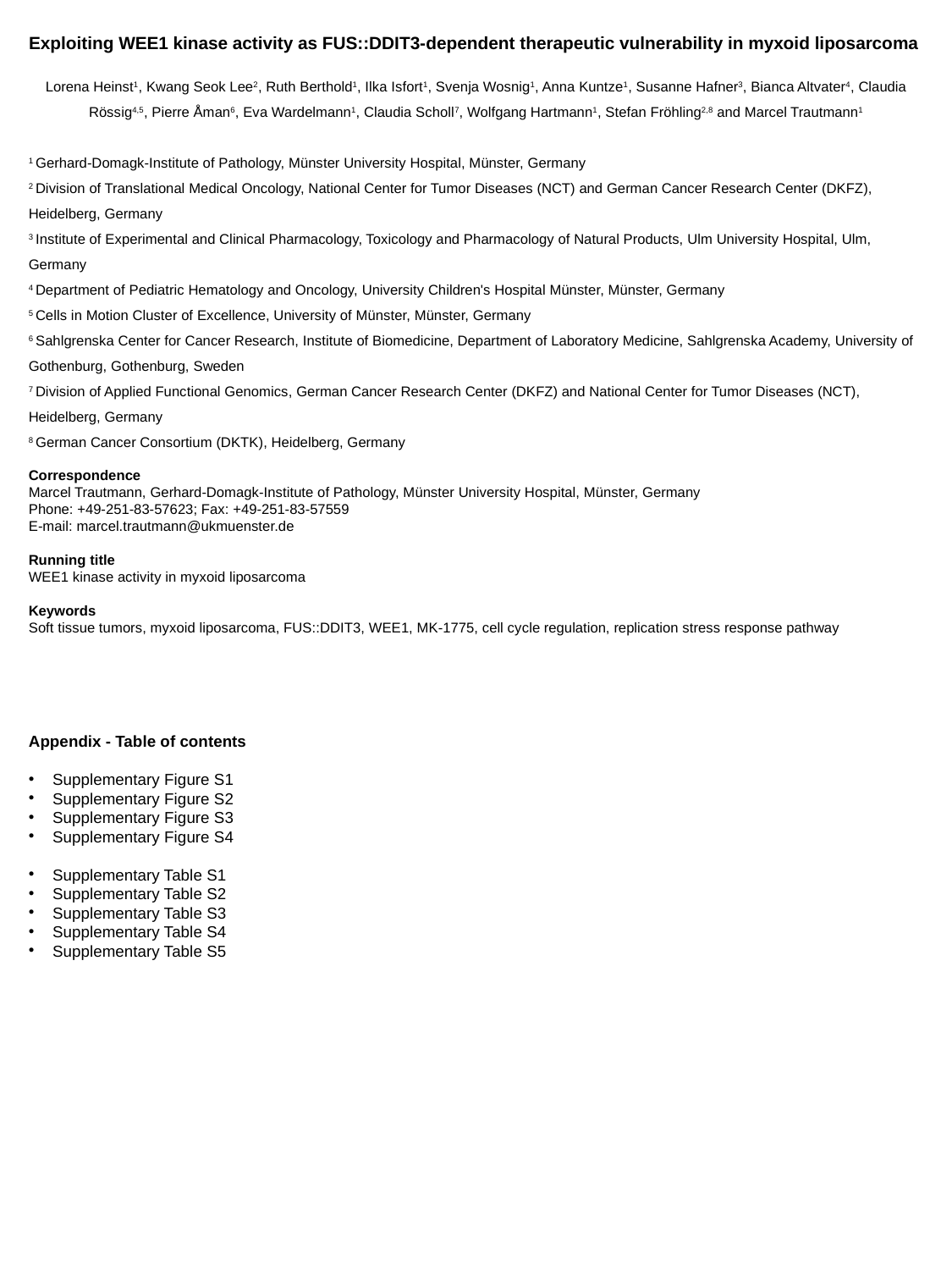

Exploiting WEE1 kinase activity as FUS::DDIT3-dependent therapeutic vulnerability in myxoid liposarcoma
Lorena Heinst^1^, Kwang Seok Lee^2^, Ruth Berthold^1^, Ilka Isfort^1^, Svenja Wosnig^1^, Anna Kuntze^1^, Susanne Hafner^3^, Bianca Altvater^4^, Claudia Rössig^4,5^, Pierre Åman^6^, Eva Wardelmann^1^, Claudia Scholl^7^, Wolfgang Hartmann^1^, Stefan Fröhling^2,8^ and Marcel Trautmann^1^
^1^ Gerhard-Domagk-Institute of Pathology, Münster University Hospital, Münster, Germany
^2^ Division of Translational Medical Oncology, National Center for Tumor Diseases (NCT) and German Cancer Research Center (DKFZ), Heidelberg, Germany
^3^ Institute of Experimental and Clinical Pharmacology, Toxicology and Pharmacology of Natural Products, Ulm University Hospital, Ulm, Germany
^4^ Department of Pediatric Hematology and Oncology, University Children's Hospital Münster, Münster, Germany
^5^ Cells in Motion Cluster of Excellence, University of Münster, Münster, Germany
^6^ Sahlgrenska Center for Cancer Research, Institute of Biomedicine, Department of Laboratory Medicine, Sahlgrenska Academy, University of Gothenburg, Gothenburg, Sweden
^7^ Division of Applied Functional Genomics, German Cancer Research Center (DKFZ) and National Center for Tumor Diseases (NCT), Heidelberg, Germany
^8^ German Cancer Consortium (DKTK), Heidelberg, Germany
Correspondence
Marcel Trautmann, Gerhard-Domagk-Institute of Pathology, Münster University Hospital, Münster, Germany

Running title
WEE1 kinase activity in myxoid liposarcoma
Keywords
Soft tissue tumors, myxoid liposarcoma, FUS::DDIT3, WEE1, MK-1775, cell cycle regulation, replication stress response pathway
Appendix - Table of contents
Supplementary Figure S1
Supplementary Figure S2
Supplementary Figure S3
Supplementary Figure S4
Supplementary Table S1
Supplementary Table S2
Supplementary Table S3
Supplementary Table S4
Supplementary Table S5

### Slide 2
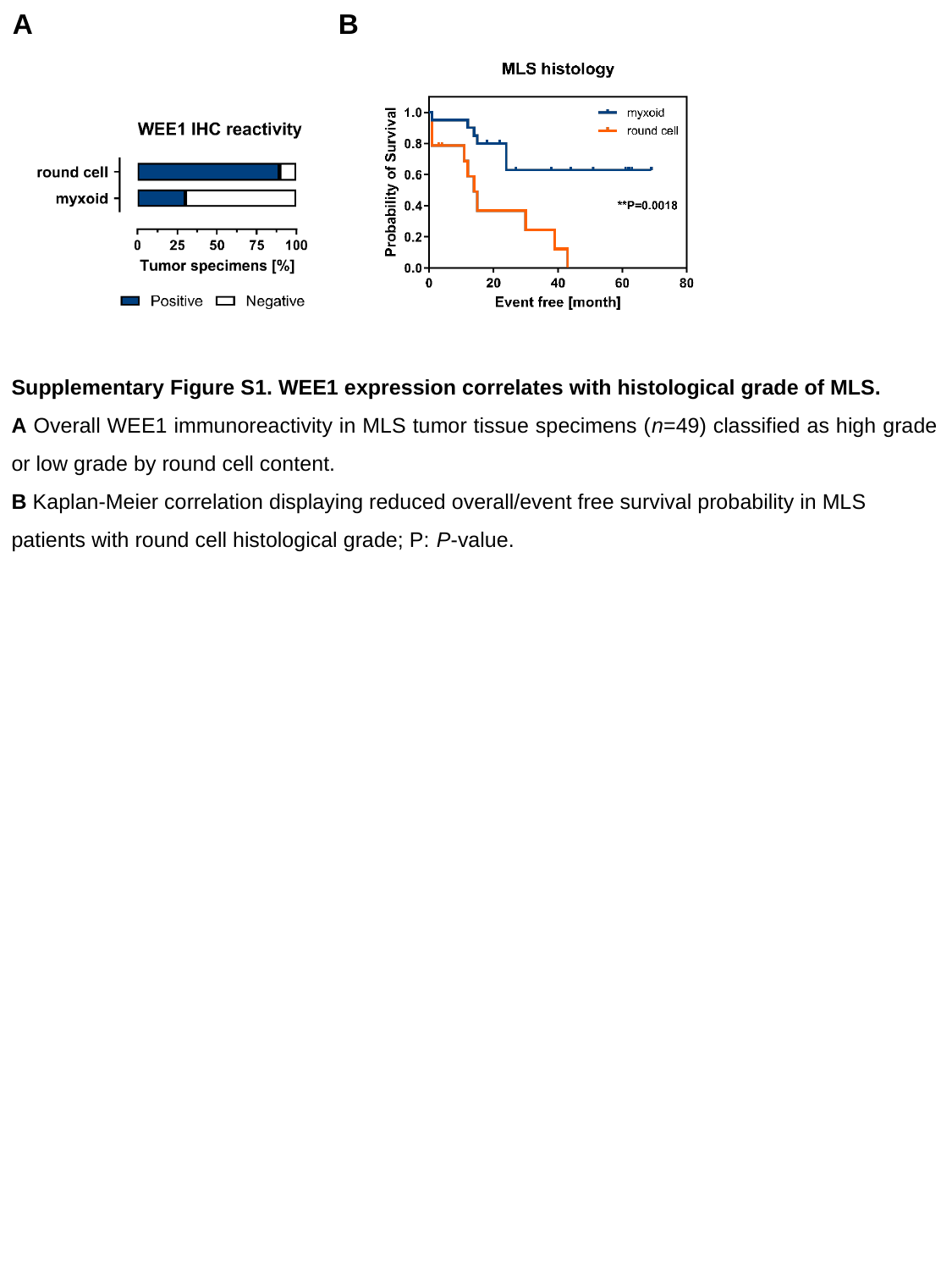

B
A
Supplementary Figure S1. WEE1 expression correlates with histological grade of MLS.
A Overall WEE1 immunoreactivity in MLS tumor tissue specimens (n=49) classified as high grade or low grade by round cell content.
B Kaplan-Meier correlation displaying reduced overall/event free survival probability in MLS patients with round cell histological grade; P: P-value.

### Slide 3
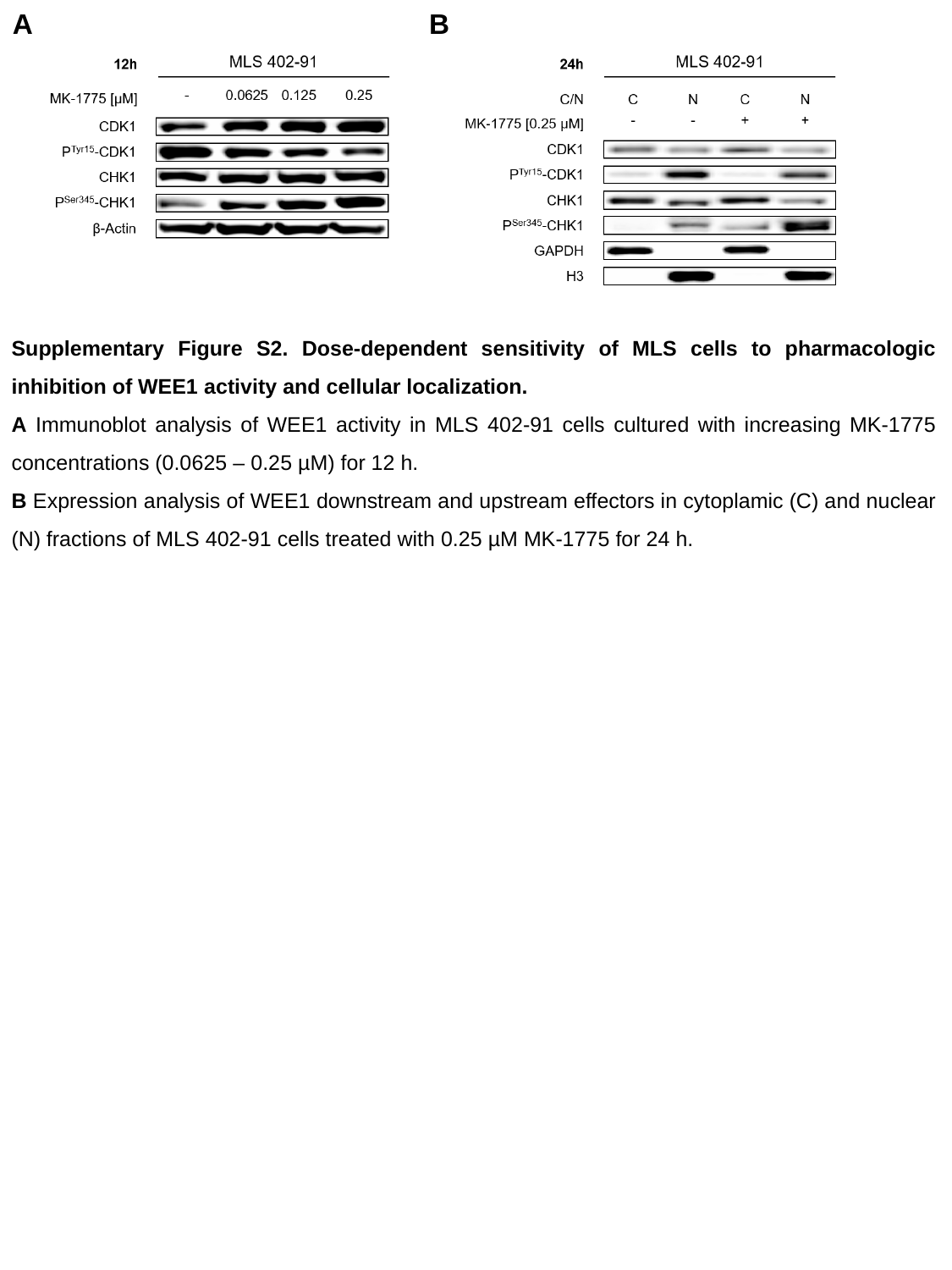

B
A
Supplementary Figure S2. Dose-dependent sensitivity of MLS cells to pharmacologic inhibition of WEE1 activity and cellular localization.
A Immunoblot analysis of WEE1 activity in MLS 402-91 cells cultured with increasing MK-1775 concentrations (0.0625 – 0.25 µM) for 12 h.
B Expression analysis of WEE1 downstream and upstream effectors in cytoplamic (C) and nuclear (N) fractions of MLS 402-91 cells treated with 0.25 µM MK-1775 for 24 h.

### Slide 4
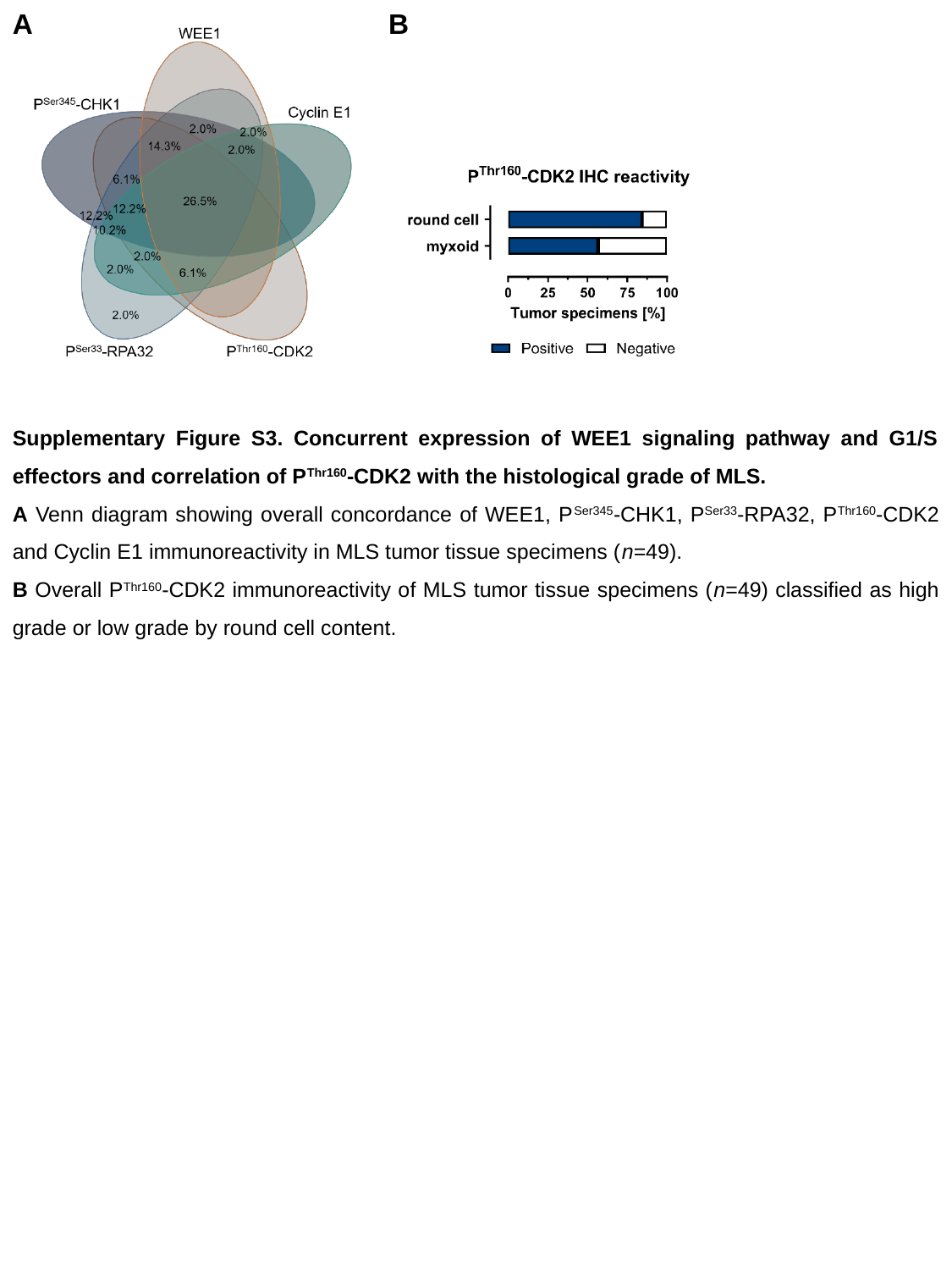

B
A
Supplementary Figure S3. Concurrent expression of WEE1 signaling pathway and G1/S effectors and correlation of P^Thr160^-CDK2 with the histological grade of MLS.
A Venn diagram showing overall concordance of WEE1, P^Ser345^-CHK1, P^Ser33^-RPA32, P^Thr160^-CDK2 and Cyclin E1 immunoreactivity in MLS tumor tissue specimens (n=49).
B Overall P^Thr160^-CDK2 immunoreactivity of MLS tumor tissue specimens (n=49) classified as high grade or low grade by round cell content.

### Slide 5
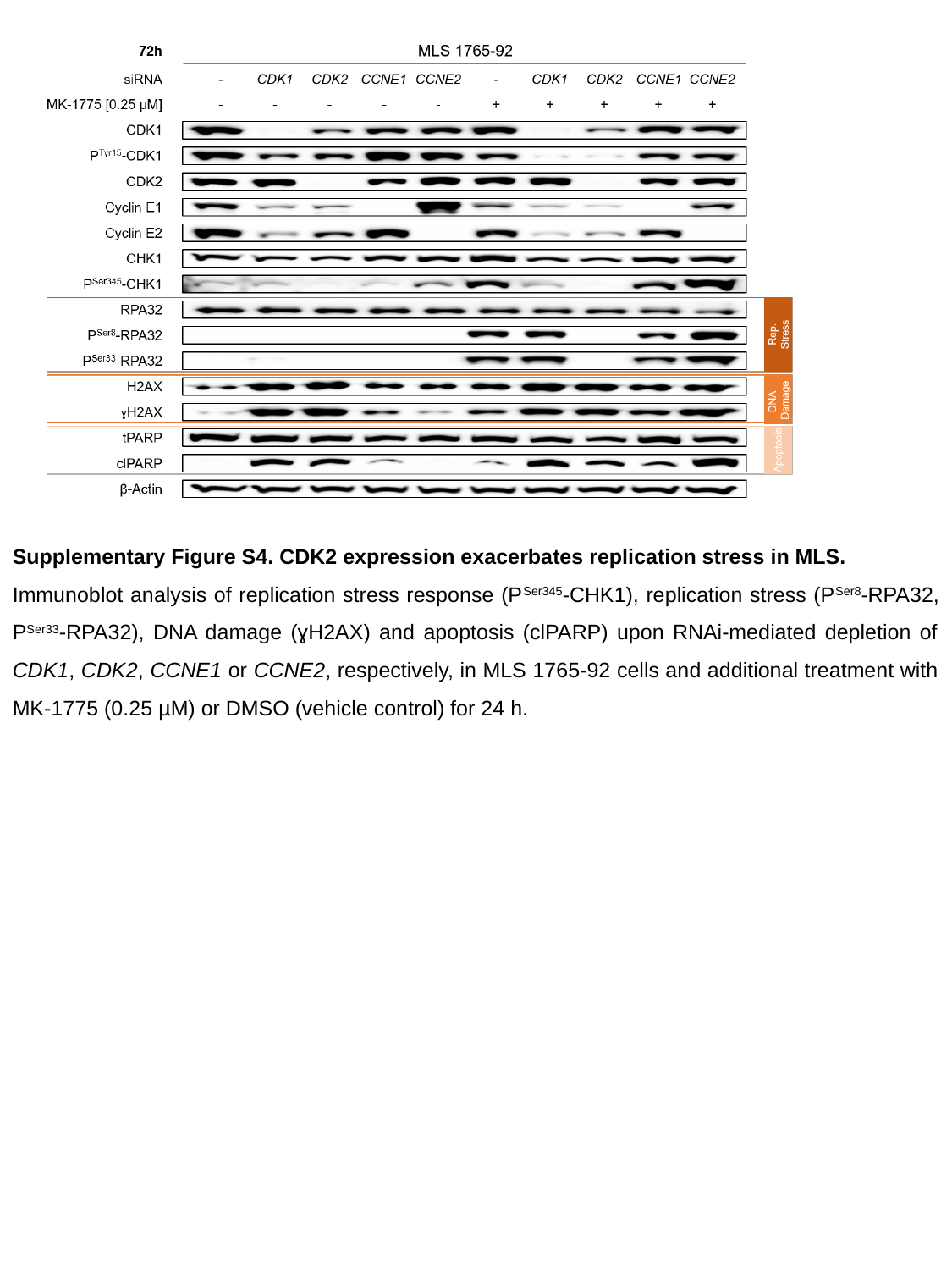

Supplementary Figure S4. CDK2 expression exacerbates replication stress in MLS.
Immunoblot analysis of replication stress response (P^Ser345^-CHK1), replication stress (P^Ser8^-RPA32, P^Ser33^-RPA32), DNA damage (ɣH2AX) and apoptosis (clPARP) upon RNAi-mediated depletion of CDK1, CDK2, CCNE1 or CCNE2, respectively, in MLS 1765-92 cells and additional treatment with MK-1775 (0.25 µM) or DMSO (vehicle control) for 24 h.

### Slide 6
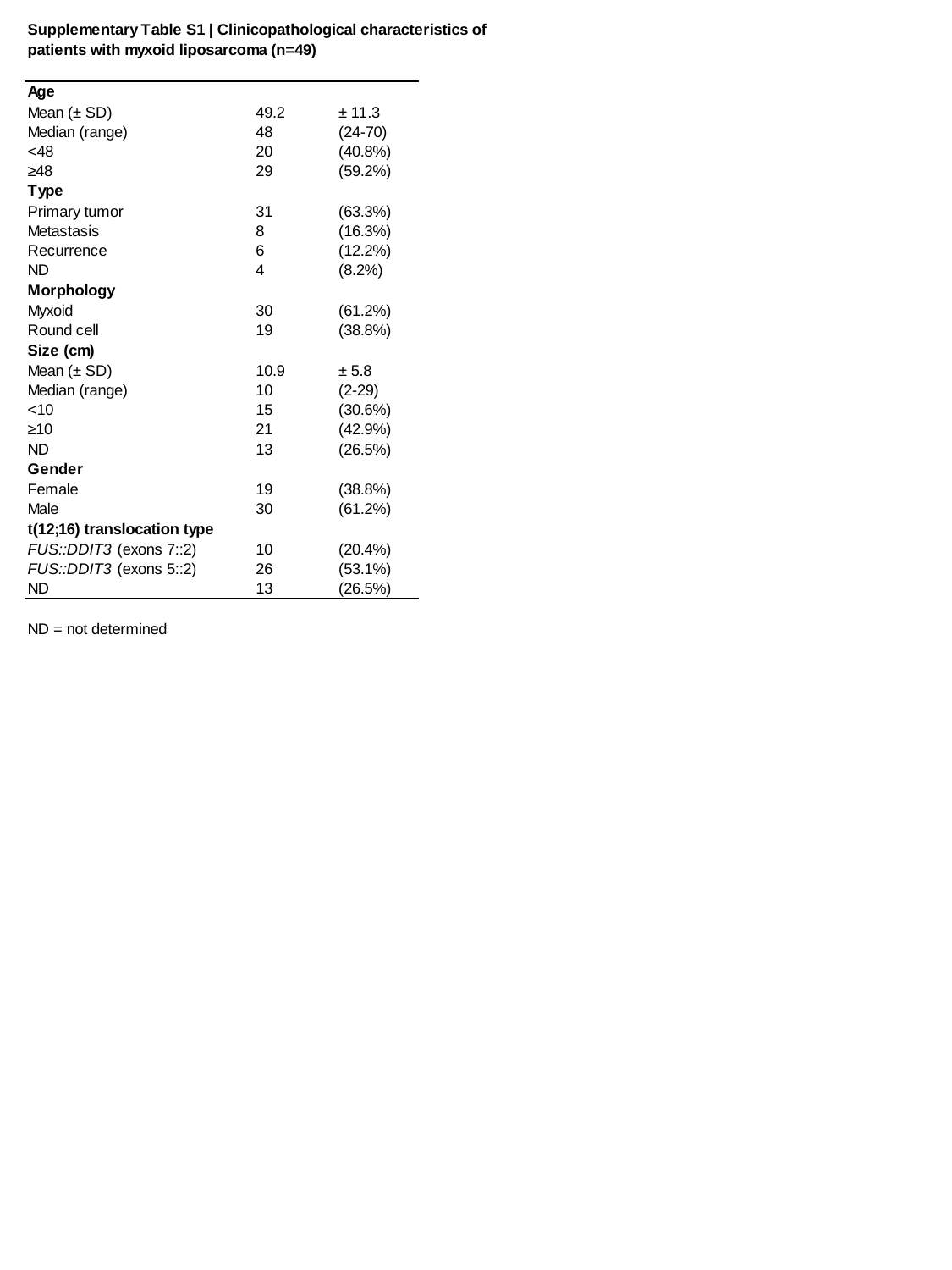

### Slide 7
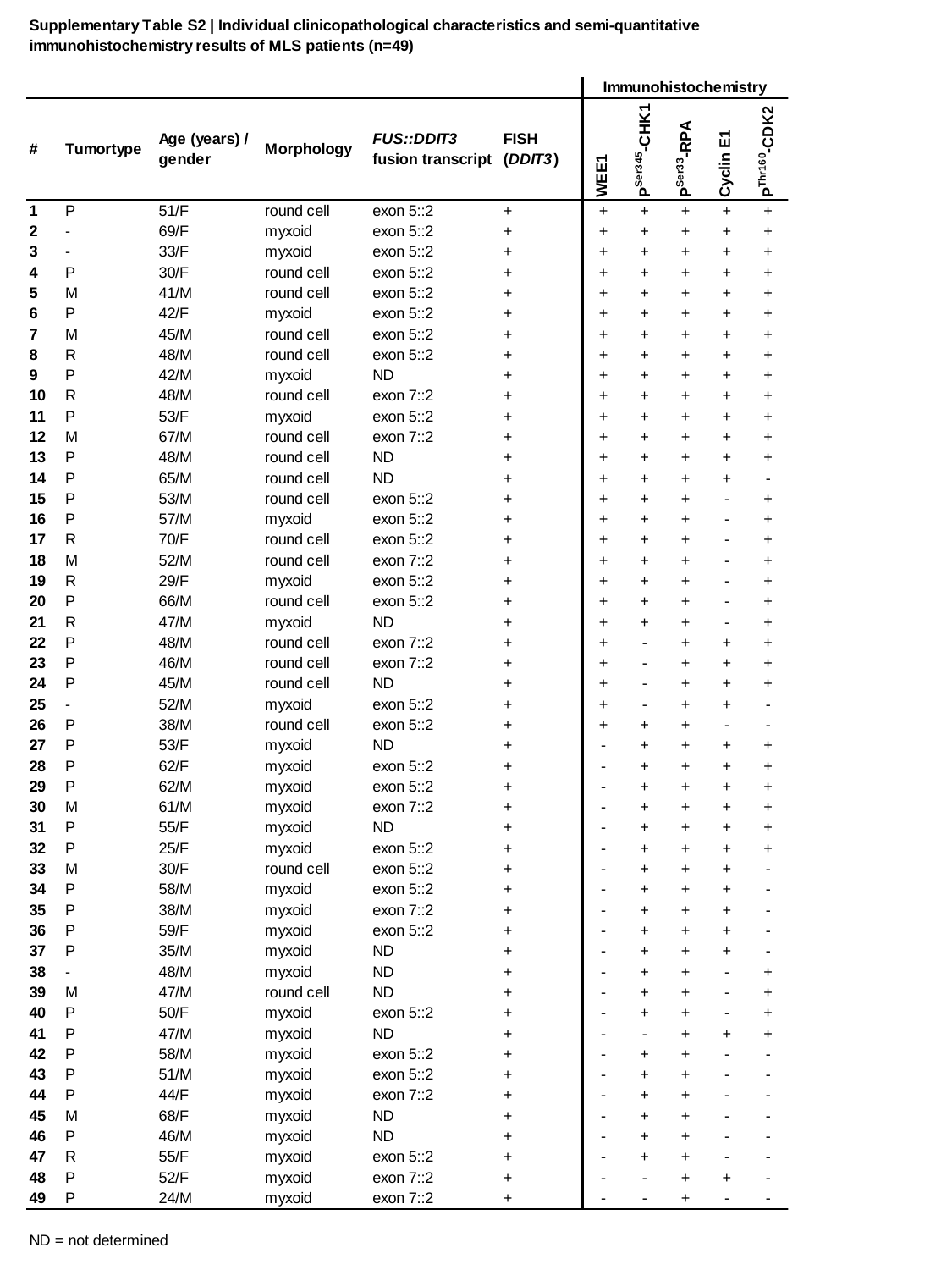

### Slide 8
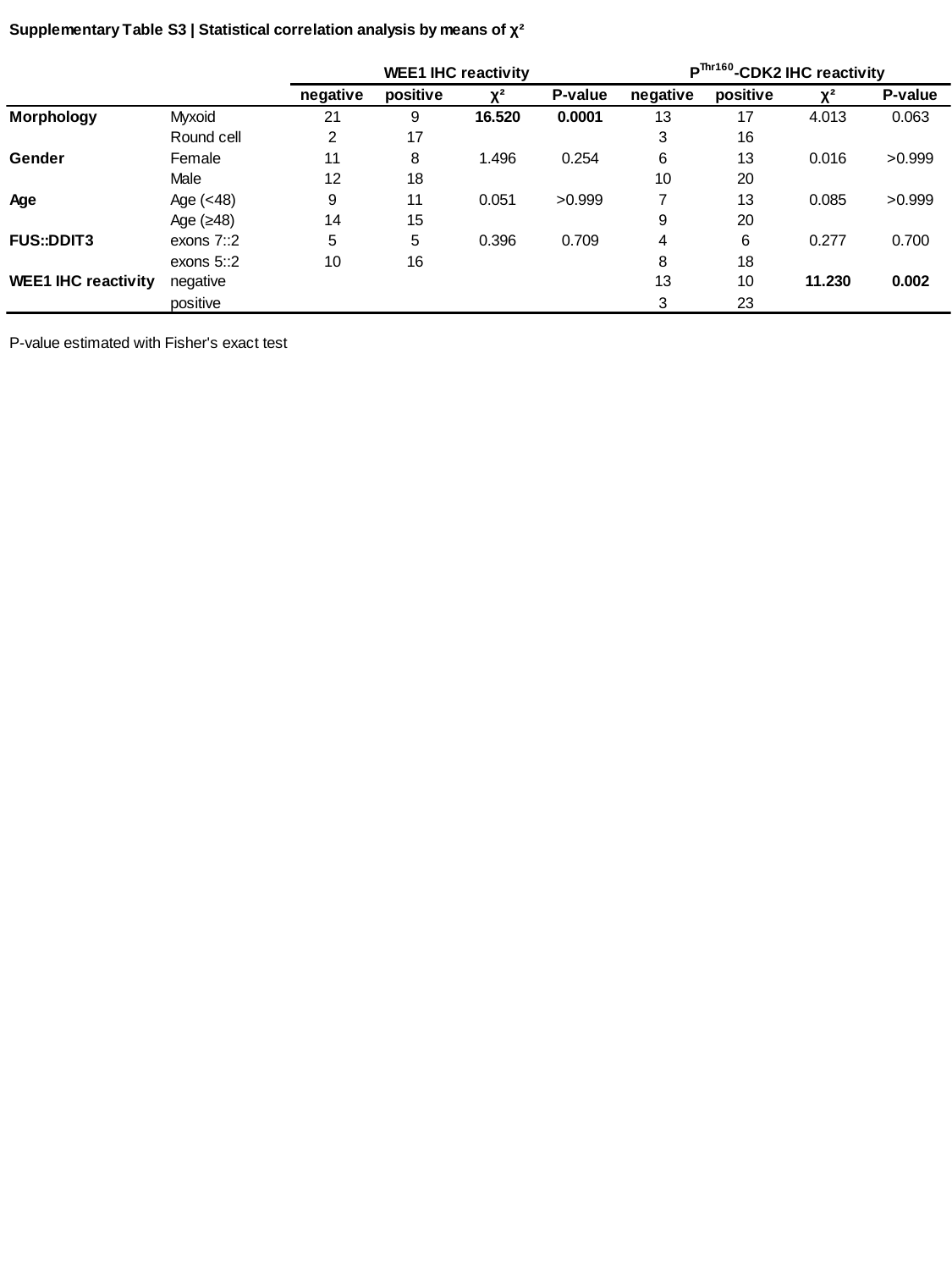

### Slide 9
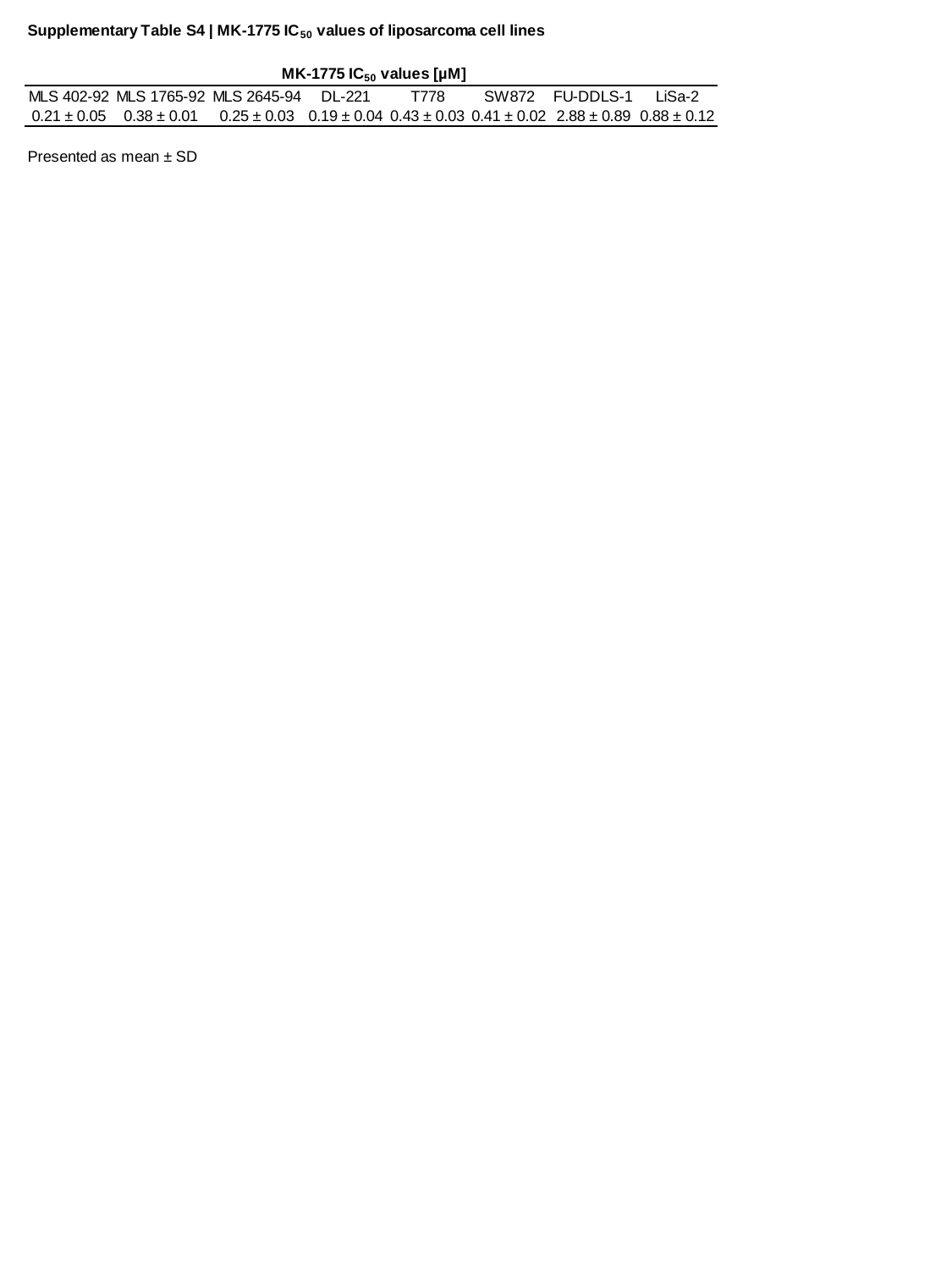

### Slide 10
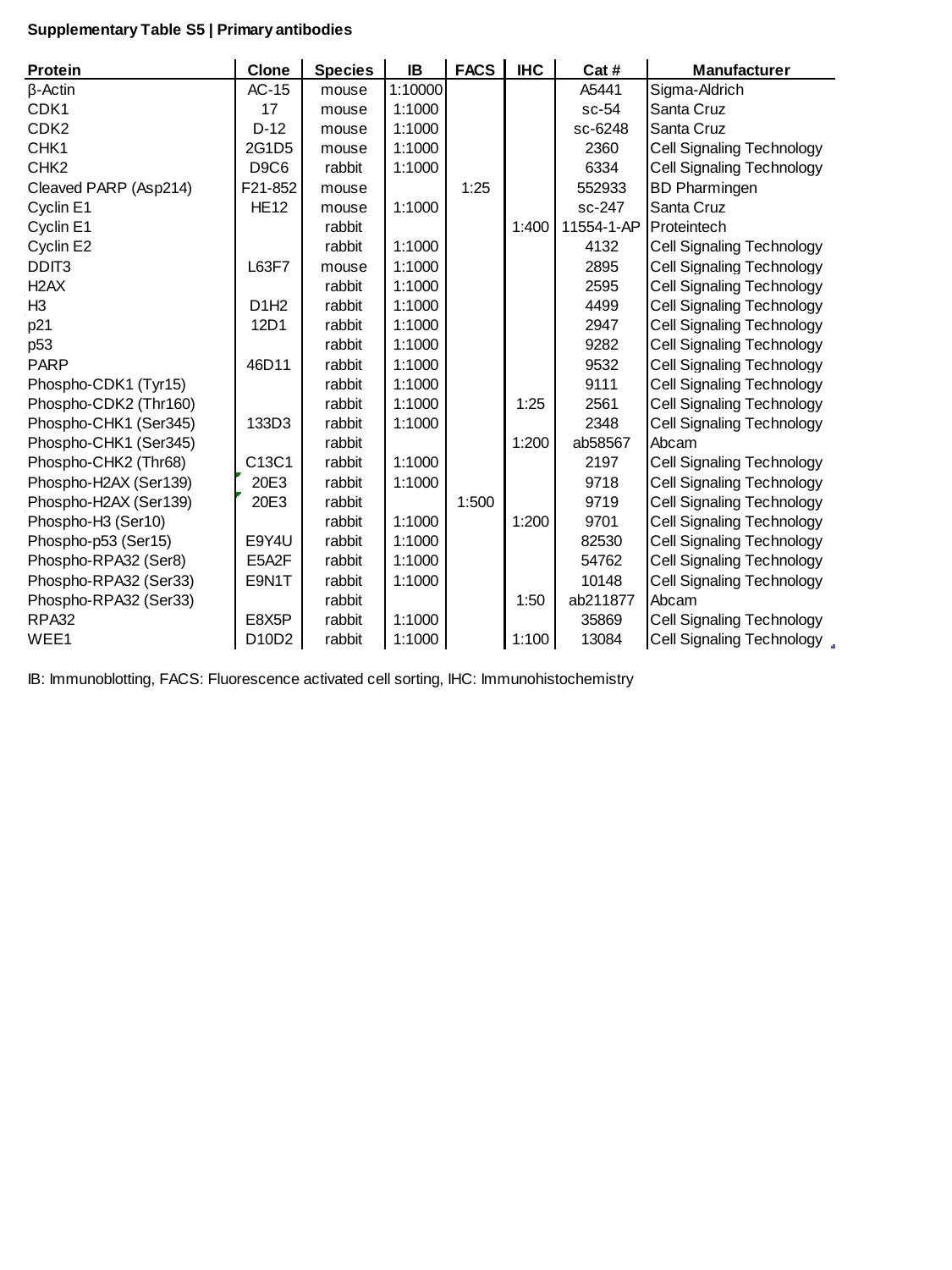
